# Games of risk and reward in carnivore communities

**DOI:** 10.1101/2021.01.27.428481

**Authors:** Joel Ruprecht, Charlotte E. Eriksson, Tavis D. Forrester, Derek B. Spitz, Darren A. Clark, Michael J. Wisdom, Marcus Bianco, Mary M. Rowland, Joshua B. Smith, Bruce K. Johnson, Taal Levi

**Affiliations:** Department of Fisheries and Wildlife, Oregon State University; Oregon Department of Fish and Wildlife; Environmental Studies Department, University of California, Santa Cruz; USDA Forest Service Pacific Northwest Research Station

**Keywords:** Carnivore, facilitation, mesopredator release, scavenging, suppression

## Abstract

Mesopredator release theory suggests that dominant predators suppress subordinate carnivores and ultimately shape community dynamics, but the assumption that subordinate species are only negatively affected ignores the possibility of facilitation through scavenging. We examined the interplay within a carnivore community consisting of cougars, coyotes, black bears, and bobcats using contemporaneous Global Positioning System telemetry data from 51 individuals, diet analysis from 972 DNA-metabarcoded scats, and data from 128 physical investigations of cougar kill sites, 28 of which were monitored with remote cameras. Resource provisioning from competitively-dominant cougars to coyotes through scavenging was so prolific as to be an overwhelming determinant of coyote behavior, space use, and resource acquisition. This was evident via strong attraction of coyotes to cougar kill sites, frequent scavenging of cougar-killed prey, and coyote diets that nearly matched cougars in the magnitude of ungulate consumption. Yet coyotes were often killed by cougars and used space to minimize encounters, complicating the fitness benefits gained from scavenging. We estimated that 23% (95% CI: 8–55%) of the coyote population in our study area was killed by cougars annually suggesting that coyote interactions with cougars are a complex behavioral game of risk and reward. In contrast, we found no indication that bobcat space use or diet was influenced by cougars. Black bears avoided cougars, but there was no evidence of attraction to cougar kill sites, and much lower levels of ungulate consumption and carcass visitation than for coyotes. Interspecific interactions among carnivores are multifaceted encompassing both suppression and facilitation.

**Significance Statement:** An incomplete understanding of the total influence competitively-dominant predators exert on subordinate species hinders our ability to anticipate the effects that changing carnivore populations will have in ecological communities. Here we show that cougars are the architects of a complex behavioral game of risk and reward, as subordinate or co-occurring carnivores are both provisioned and preyed on by the dominant predators. Each co-occurring carnivore species considered here employed a different strategy to approach the risk-reward tradeoff suggesting there are multiple viable solutions to the game. By not considering the multitude of effects dominant predators have on other carnivores, we are missing important linkages in terrestrial food webs.

## Introduction

The mesopredator release hypothesis posits a trophic cascade wherein larger-bodied and competitively-dominant predators suppress subordinate predators with consequences for prey populations (1–4). Such suppression is often the result of interspecific killing that may be so pervasive as to account for one- to two-thirds of the total mortality of terrestrial carnivores (5–7). Subordinate species may respond to this risk by altering their space use or activity patterns to minimize the probability of encounters with dominant predators (8, 9), but nonetheless, the antagonism between predator species can result in reductions in density or complete exclusion of subordinate carnivores from certain habitats or regions (4, 10, 11).

The assumption that only negative interactions influence carnivore community structure is likely overly simplistic because scavenging of dominant predator kills can also provision subordinate species (7, 12, 13). While the prey killed by dominant predators may represent a ‘fatal attraction’ that leads mesopredators to their death, the energetic rewards from scavenging in some cases may outweigh the risks (14). For example, if the risk of scavenging is the same as hunting alternative prey but yields a higher probability of finding food, the fitness-maximizing decision for the subordinate or co-occurring species should be to scavenge (Fig. 1; *SI Appendix*, Text S1). But as risks associated with scavenging increase, the optimal choice becomes a more nuanced function of the relative risk versus reward and depends on the energetic state of the potential scavenger (Fig. 1a; *SI Appendix*, Text S1). And, of course, the scavenger must make this assessment under imperfect knowledge of the risk involved. Thus, the decision for a subordinate species to scavenge from a dominant species represents a complex behavioral game of risk and reward (Fig. 1b). How the costs and benefits of interactions with dominant predators influence the overall fitness of subordinate or co-occurring species has rarely been examined.

**Figure 1.**
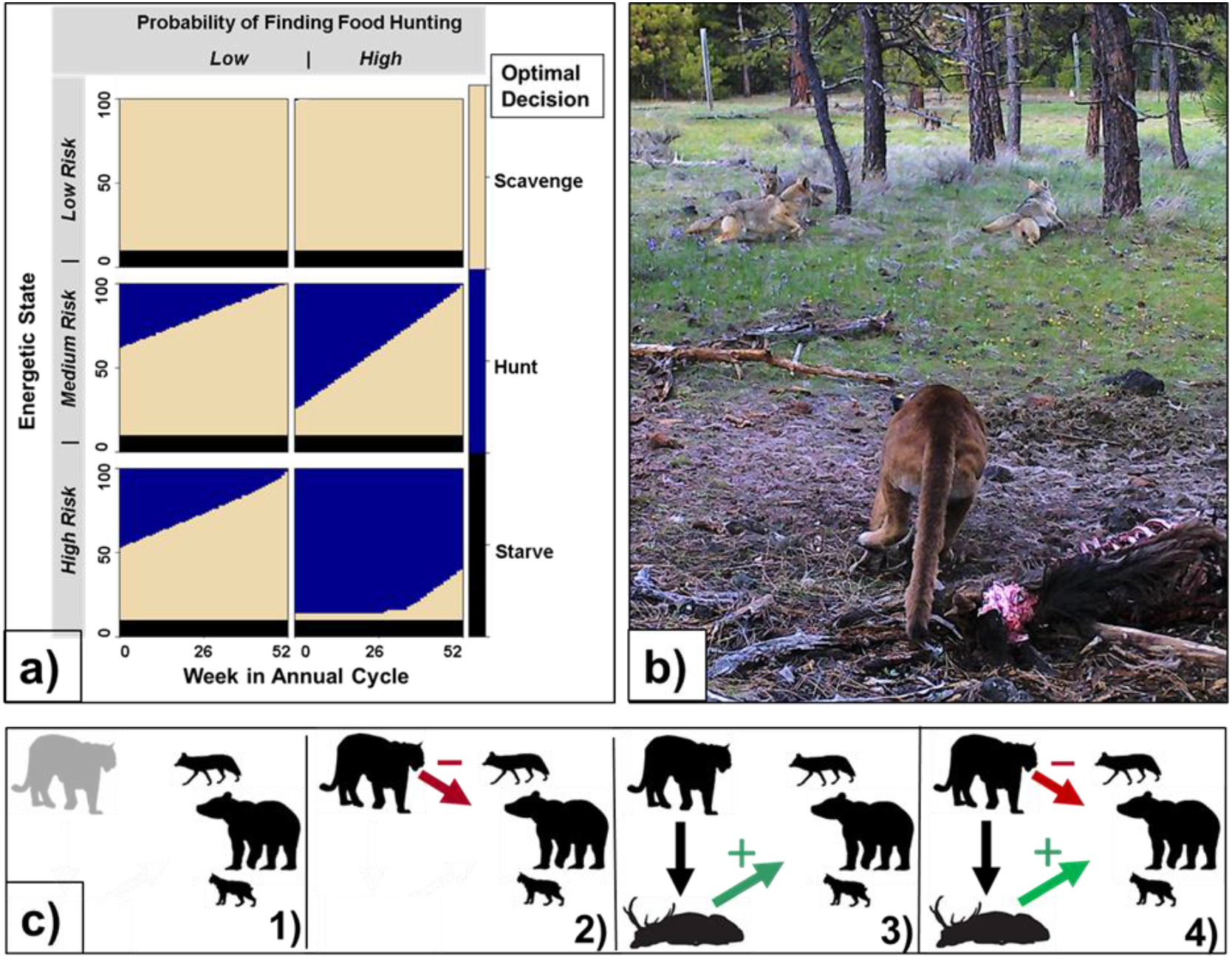
a) Predictions from a two patch dynamic state variable model used as an illustrative example. The conceptual model predicts the fitness-maximizing decision where the choices are to scavenge (patch 1, tan shading) or forego the scavenging opportunity and hunt alternative prey (patch 2, blue shading). The optimal decision in this theoretical model depends on the risk, quantified as daily probability of death, and reward, quantified as probability of finding food. The fitness-maximizing decision depends not only on risk and reward of each patch, but also the energetic state of the scavenger (y-axis) and the week within an annual cycle (x-axis). b) Interaction between coyotes and a cougar at a kill site monitored by remote camera. c) Possible influences of dominant predators in a carnivore community: 1) dominant predators do not influence other carnivores (null hypothesis), 2) dominant predators negatively influence other carnivores through interference competition or interspecific killing (suppression hypothesis), 3) dominant predators positively influence other carnivores due to resource provisioning via scavenging (facilitation hypothesis), and 4) dominant predators both positively and negatively influence other carnivores (provision-predation hypothesis if positive and negative interactions are independent, fatal attraction hypothesis if scavenging leads to increased mortality).

Understanding the intraguild dynamics of carnivores is critical to anticipating the ecosystem-level consequences of changing predator populations, but establishing the relative prevalence of suppression and facilitation among carnivores has remained intractable. Strong inference about the magnitude and direction of behavioral and population-level interactions among carnivores requires (1) quantifying the intraguild predation rate on subordinate or co-occurring carnivores, (2) understanding the extent to which subordinate or co-occurring carnivores utilize carrion from dominant predator kills, and (3) evaluating whether dominant predators or their kills influence the movements or space use of subordinate or co-occurring carnivores. Spatial avoidance of dominant predators and high rates of intraguild predation would suggest other carnivores are suppressed. By contrast, high utilization of dominant predator kills by subordinates would support the facilitation hypothesis. We note that while the literature often refers to the benefit conferred by scavenging as facilitation (7, 12, 14), this term implies no harm to the dominant species. If carrion is lost due to scavenging while the owner of the kill is still utilizing the carcass (15, 16), the phenomenon may be more appropriately termed parasitism, or more specifically, kleptoparasitism.

Here we quantify the influence of cougars (*Puma concolor*) on coyotes (*Canis latrans*), black bears (*Ursus americanus*), and bobcats (*Lynx rufus*). Cougars are widely considered dominant to coyotes and bobcats, and although previous researchers have suggested cougars are subordinate to black bears, the available literature suggests that cougars kill bears more often than the reverse (17). We therefore consider bears and cougars to be co-occurring and not necessarily fitting within a clear dominance hierarchy. We leveraged four empirical datasets including contemporaneous Global Positioning System (GPS) telemetry data of 51 individuals across the four species of carnivores, diet analysis from 972 DNA-metabarcoded carnivore scats, and data from 128 physical investigations of cougar kill sites, 28 of which were monitored for scavenging via remote cameras to evaluate how sympatric carnivores are influenced by cougars. We hypothesized that black bears, bobcats and coyotes could be unaffected by cougars (null hypothesis), negatively influenced by cougars (suppression hypothesis), positively influenced by cougars (facilitation hypothesis), or both positively and negatively affected by cougars (“provision-predation hypothesis” if the positive and negative interactions are independent, “fatal attraction hypothesis” if scavenging leads to increased mortality) (Fig. 1c). We used contemporaneous GPS data to infer whether each carnivore species altered movements due to the proximity of a cougar or cougar kill site. We hypothesized that carnivore species attracted to cougar kill sites would have a high frequency of carrion in their diet—suggesting facilitation from cougars—but would also be killed by cougars at higher rates than species that avoided cougar kill sites, indicating suppression. Together, these approaches provided a robust evaluation of the fine-scale behavioral interactions and facilitation versus suppression tradeoff among species within carnivore communities, which will be increasingly important as predator populations recover across portions of their former range and are extirpated in others.

## Materials and Methods

### Data collection

We collected data on four sympatric carnivore species (cougars, coyotes, black bears, and bobcats) in and adjacent to the Starkey Experimental Forest and Range in the Blue Mountains of northeastern Oregon, USA (Fig. 2 and *SI Appendix*, Text S2) between 2016 and 2020. We tagged 17 cougars (8M, 9F), 17 coyotes (10M, 7F), 11 black bears (7M, 4F), and 6 bobcats (3M, 3F) with GPS collars (Vectronic Aerospace or Lotek Wireless, Inc.) programmed to record positions every 2 or 3 hours (*SI Appendix*, Text S3). We used conservation detection dogs (18) to locate scats (*SI Appendix*, Text S4) and genetically confirmed 774 coyote scats, 85 black bear scats, 96 bobcat scats, and 17 cougar scats (19) that were DNA-metabarcoded for diet analysis (*SI Appendix*, Text S5). We considered the frequency of occurrence of elk [*Cervus canadensis*]) in scats of subordinate carnivores as a qualitative but imperfect proxy for scavenging (14). It is unlikely any of the subordinate carnivores could kill adult elk so any elk found in scats must be from scavenging or direct predation on calves. However, elk consumption by coyotes and bobcats is likely exclusively from scavenging because these species rarely kill elk calves in the region (20). Nonetheless, the frequency of occurrence of elk in scats, particularly for bears, can only be interpreted as the maximum possibly attained from scavenging because some unknown fraction may have come from predation of neonates. In addition to inference from scat analysis, we further quantified cougar diets by conducting ground searches of clusters of cougar GPS locations indicating potential kill sites of tagged individuals (*SI Appendix*, Text S6) (21). At 28 of the 128 confirmed cougar kill sites, we deployed remote cameras (Bushnell Trophy Cam Aggressor, Browning Dark Ops HD Pro, and Reconyx HyperFire 2) to estimate scavenging rates by the other carnivores. We separately tallied the number of daily visits made by coyotes, bears and bobcats to cougar kill sites that were detected on camera, where a visit was defined as a detection separated by at least 30 minutes from another detection of a given species. We used negative binomial regression to model the number of daily visits to the carcass by a given species as:

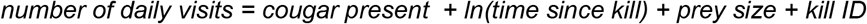

where cougar present (1/0) indicates whether the cougar was present at the kill that day, time since kill is the number of days since the kill was made, prey size (1/0) indicates whether the prey was large (adult ungulate) or not (young of the year ungulate), and kill ID is a random intercept identifying a given kill site.

**Figure 2.**
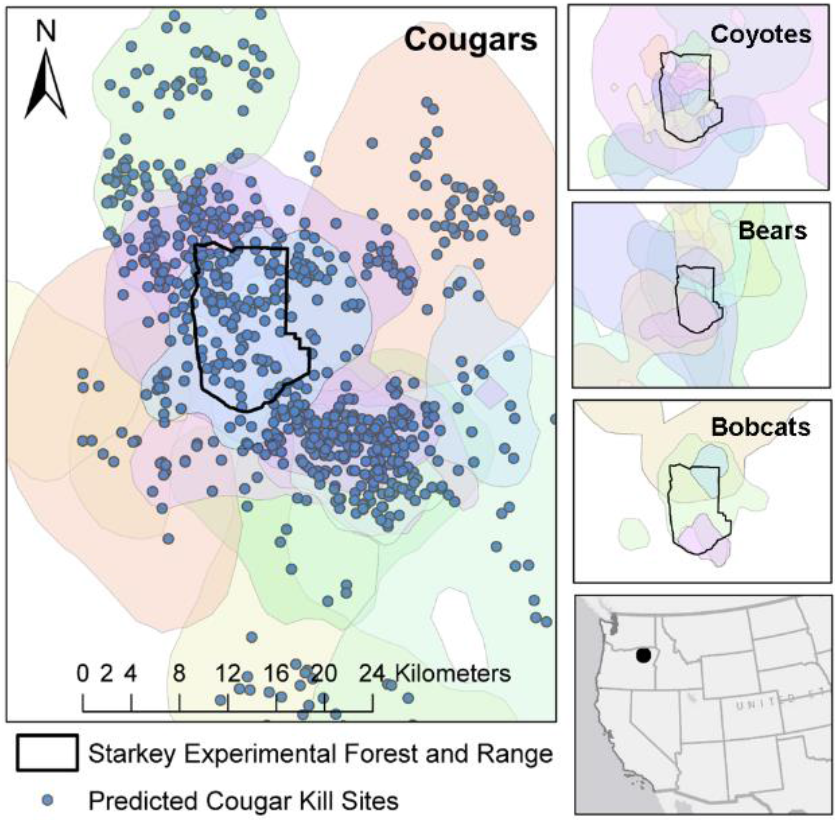
Study area. Left panel: Known and predicted cougar kill sites (blue points) and 95% kernel density home range estimates (colored polygons) for collared cougars. Right panels, top to bottom: coyote, black bear and bobcat 95% kernel density home range estimates (colored polygons). All panels: Starkey Experimental Forest and Range (black polygon). Kernel density estimates were calculated using the kernelUD function in the adehabitatHR package (53) in program R. We used the default ad hoc smoothing parameter which assumes the kernel is bivariate normal (53).

### Calculating Intraguild Predation Rate

At times, the prey items found at cougar kill sites were other carnivores which allowed us to calculate the proportion of cougar kills representing intraguild predation. Combining these data with information on contemporaneously-estimated carnivore densities (22) and cougar kill rates (21) from a recent study in the area allowed us to calculate intraguild predation rates, i.e., the proportion of the co-occurring carnivore population dying annually due to cougar predation. We calculated the intraguild predation rate on species *i* (*P_i_*) as the number of individuals from species *i* killed per unit time (*K_i_*) divided by the population density of species *i* (*D_i_*) where *K_i_* is the product of cougar population density (*D_c_*) cougar kill rate (kills/year), *R*, and proportion of kills corresponding to species *i* (*F_i_*):

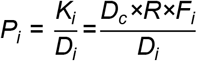

We quantified uncertainty in *P_i_* by propagating error inherent in each parameter using Markov Chain Monte Carlo simulations (*SI Appendix*, Text S7).

### Step selection functions

We fit integrated step-selection functions (hereafter *SSFs*) (23, 24) to GPS collar data to estimate how coyote, bobcat, and bear movements were influenced by landscape variables, cougar proximity, or cougar kill site proximity. We included movement attributes as parameters in each model to reduce bias arising from sampling control locations (25) using a gamma distribution to characterize the natural log of step lengths and a von Mises distribution to characterize the cosine of turning angles (24, 26). We generated 20 random steps for each observed step (27). To ensure that territoriality did not bias inference, we did not allow the endpoints of random steps to fall outside a 1,000 m buffer of the 99% minimum convex polygon of each animal’s territory. We did this to prevent detecting spurious avoidance behavior due to a random step occurring in an area for which the animal did not have uninhibited access due to territoriality.

We sought to identify whether landscape features and the proximity of a cougar (hereafter, ‘distance to cougar’ or D2C) or cougar kill site (hereafter, ‘distance to kill’ or D2K) influenced the movements of coyotes, bobcats, and bears. We calculated distance to cougar as the natural log-transformed Euclidean distance between each focal individual (coyote, bobcat, or bear) and the nearest GPS-collared cougar at that time to determine whether the subsequent movements of the animal were toward or away from the cougar. To restrict encounters to only those in which the cougar could plausibly be detected by the subordinate species, we only allowed cougars within 1,000 m of the focal bear, bobcat, or coyote to influence the movements of the other carnivores by using a binary indicator for whether a cougar was present (hereafter, ‘cougar present’ or CP) (1/0) at each GPS fix. We cast ‘distance to cougar’ as an interaction term with ‘cougar present’ such that the term was nullified when the nearest known cougar was > 1,000 m away. We used a buffer of 1,000 m because that is the distance elk alter behavior in response to wolves (*Canis lupus*) (28) and has also been used to determine interaction distances between cheetahs (29). Further, it is implausible that species could detect cougars at distances greatly exceeding this value. However, to ensure this cutoff did not change our inference about the direction or magnitude of the effect of cougars, we replicated the analyses using additional cutoff values of 500 and 1,500 m. If GPS fixes between cougars and the other carnivores were not contemporaneous (i.e. falling on the same hour plus a tolerance of 300 seconds) due to misalignment or missed fixes, we imputed the location of the cougar using a correlated random walk (30). We allowed cougar locations to be imputed if a single GPS fix was missed but we censored location data for any gaps exceeding 6 hours.

We used this same approach to assess how carnivores responded to cougar kill sites by including a term for ‘distance to kill,’ calculated as the natural log of the Euclidean distance between each GPS position of a focal individual and the nearest cougar kill site. Potential kill sites were identified from clusters of sequential cougar GPS locations indicating reduced movement and were either confirmed via physical investigations or identified as probable kills using a predictive model (21) (*SI Appendix*, Text S6). We paired ‘distance to kill’ with a binary indicator variable describing whether there was a known or predicted cougar kill present (hereafter, ‘kill present’ or KP) (1/0) within a 3,000 m buffer of the focal individual. When there was not a kill within the buffer, the indicator variable took the value of zero and the whole term became null. If there was more than one kill within this buffer for a given individual at a given time, we considered only the distance to the most recent kill. Our assumption that carnivores could detect a carcass up to distances of 3,000 m was conservative; for example, one GPS-collared coyote in our study exhibited two bouts of directed travel of 2.2 km and 3.6 km outside its territory to feed on an elk carcass suggesting an ability to detect carrion from these distances (*SI Appendix*, Fig. S1). Further, previous studies report that coyotes can travel between 12.2 and 20.5 km to reach carcasses of domestic animals (31, 32), while arctic foxes (*Vulpes lagopus*) routinely detect marine mammal carcasses at distances of at least 10 km (33). While less is known about olfaction distances in bears, it is reported that polar bears (*Ursus maritimus*) can detect seal breathing holes at distances of 3 km (34). While the available literature suggests 3,000 m is an appropriate value for a cutoff, we replicated each analysis using cutoffs of 2,000 m and 4,000 m to ensure inference was not an artifact of the buffer chosen.

To avoid data contamination due to old carcasses that no longer provided scavenging opportunities, we classified all kills occurring within 30 days of the focal fix as present by censoring kills exceeding this threshold. To quantify how movements of carnivores toward or away from the kill may change as the carcass aged, we created another variable cast as an interaction between the kill present indicator variable (KP), the natural log of the distance to kill (D2K), and the natural log of the time elapsed since the kill (hereafter, ‘time since kill’ or TSK). This interaction allowed the magnitude of avoidance or attraction to change as a function of the age of the carcass.

It is certain that at times additional cougars were present in the study area not monitored with GPS collars, so the inferences made about effects of cougars and their kill sites on the other carnivore species should be considered conservative, at least if the collared sample was representative of the entire population. Thus, with complete information on all cougar and kill site locations, the D2C and D2K variables may have had larger effect sizes and/or smaller standard errors than reported here, assuming carnivores exhibited similar behavior toward collared and uncollared cougars.

Before including cougar variables, we first fit models containing only landscape, and movement variable (hereafter, *habitat model*) according to

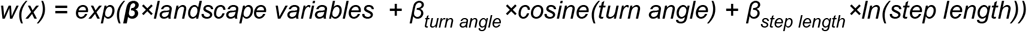

where ‘landscape variables’ refer to the continuous variables *ln*(distance to open road), *ln*(distance to perennial water source), canopy cover, topographic ruggedness, and the factor variable potential vegetation type. *A priori*, we expected each of these landscape variables to influence resource selection of each of the species considered, so for simplicity we did not conduct model selection on subsets of these variables and instead retained them all.

We then added the cougar and kill site terms and used Akaike’s Information Criterion (AIC) (35) to assess whether this full model (hereafter, *habitat + cougar model*) outperformed the habitat-only model:

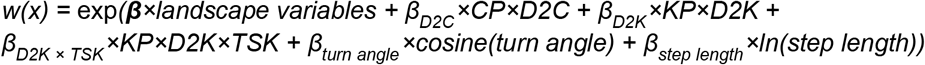

By setting *β_D2K_* × KP × D2K + *β_D2K × TSK_* × KP × D2K × TSK equal to zero and solving for TSK, we estimated the time until the effect of the kill becomes null. Assuming a kill site is initially a source of attraction that diminishes in time, this is estimated by

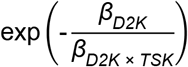

We used the ‘amt’ package (26) in Program R 3.6.1 (R Development Core Team 2019) to format data, generate random steps, and fit models.

### Evaluating Weight of Evidence for Facilitation, Suppression, and Null Hypotheses

We define suppression as the outcome of negative interactions, possibly including death or simply instilling fear to a degree that motivates changes in behavior or habitat use (36). Correspondingly, we define facilitation as the outcome of beneficial or positive interactions, including increased foraging opportunities that may improve fitness, and may motivate changes in behavior or habitat use. We leveraged multiple independent datasets on the four carnivore species using a weight of evidence approach to determine whether cougars suppress and/or facilitate black bears, bobcats, and coyotes. *A priori*, we established the following criteria: 1) spatial avoidance of cougars would be evidence of suppression, attraction to cougars would be evidence of facilitation, and indifference toward cougars would support the null hypothesis, 2) spatial avoidance of cougar kill sites would be evidence of suppression, attraction to kill sites would be evidence of facilitation, and indifference toward kill sites would support the null hypothesis, 3) a higher frequency of ungulates found in scats of subordinate or co-occurring predators (a proxy for scavenging in our system) in similar proportions to those observed in cougar scats would suggest a higher degree of facilitation than would a lower frequency of ungulates found in scats, and no ungulates found in scats would support the null hypothesis, 4) higher scavenging rates would suggest a greater degree of facilitation than would lower scavenging rates, and the absence of scavenging would support the null hypothesis, and 5) any amount of intraguild predation by cougars on other carnivores would be evidence of suppression (regardless of whether the mortality is compensatory), whereas complete absence of intraguild predation would support the null hypothesis.

## Results

### Step selection functions

Coyotes, black bears, and bobcats each responded to a subset of landscape variables; most notably, bears and bobcats strongly selected for high canopy cover while coyotes avoided high canopy cover (Table 1 and *SI Appendix*; Fig. S2). Including the three cougar parameters (CP × D2C, KP × D2K, and KP × D2K × TSK) substantially improved model fit for bears (ΔAIC = 10.36) and particularly for coyotes (ΔAIC = 23.96), but the habitat-only model outperformed the habitat + cougar model for bobcats (ΔAIC = 2.69) (*SI Appendix*, Table S1). Notably, coyotes exhibited a strong attraction to cougar kill sites (β = −
0.26, p < 0.001) which attenuated as the carcass aged (β = 0.071, p < 0.001; Fig. 3 and Table 1) and resulted in approximately 39 days of selection for cougar kill sites. Post-hoc analyses revealed that attraction of coyotes to cougar kill sites was driven primarily by resident male coyotes (*SI Appendix*, Text S9–S10, Table S2–S3). Coyotes avoided cougars (β = 0.20, p = 0.014; Table 1), but there was no evidence to suggest attraction to kill sites was diminished if a cougar was present at the kill, indicating coyotes may disregard the risk of cougars when a food reward is present (*SI Appendix*, Text S11, Table S4). Bears were indifferent to kill sites (β = 0.01, p = 0.93) but actively avoided cougars (β = 0.55, p ≤ 0.001; Table 1). Bobcats were indifferent to both cougars (β = 0.26, p = 0.33) and their kill sites (β = 0.11, p = 0.54; Table 1). A post-hoc analysis did not provide strong evidence that bobcats avoided coyotes in space and therefore coyote presence was probably not the primary reason they did not utilize cougar kills (*SI Appendix*, Text S12, Table S5). For all species, the choice of buffer for determining whether a cougar (CP) or kill (KP) was present did not appreciably change inference about the direction or magnitude of these effects except that coyote avoidance of cougars was only evident when buffers 1,000 m or less were used (*SI Appendix*, Text S13, Table S6–S11).

**Table 1.**
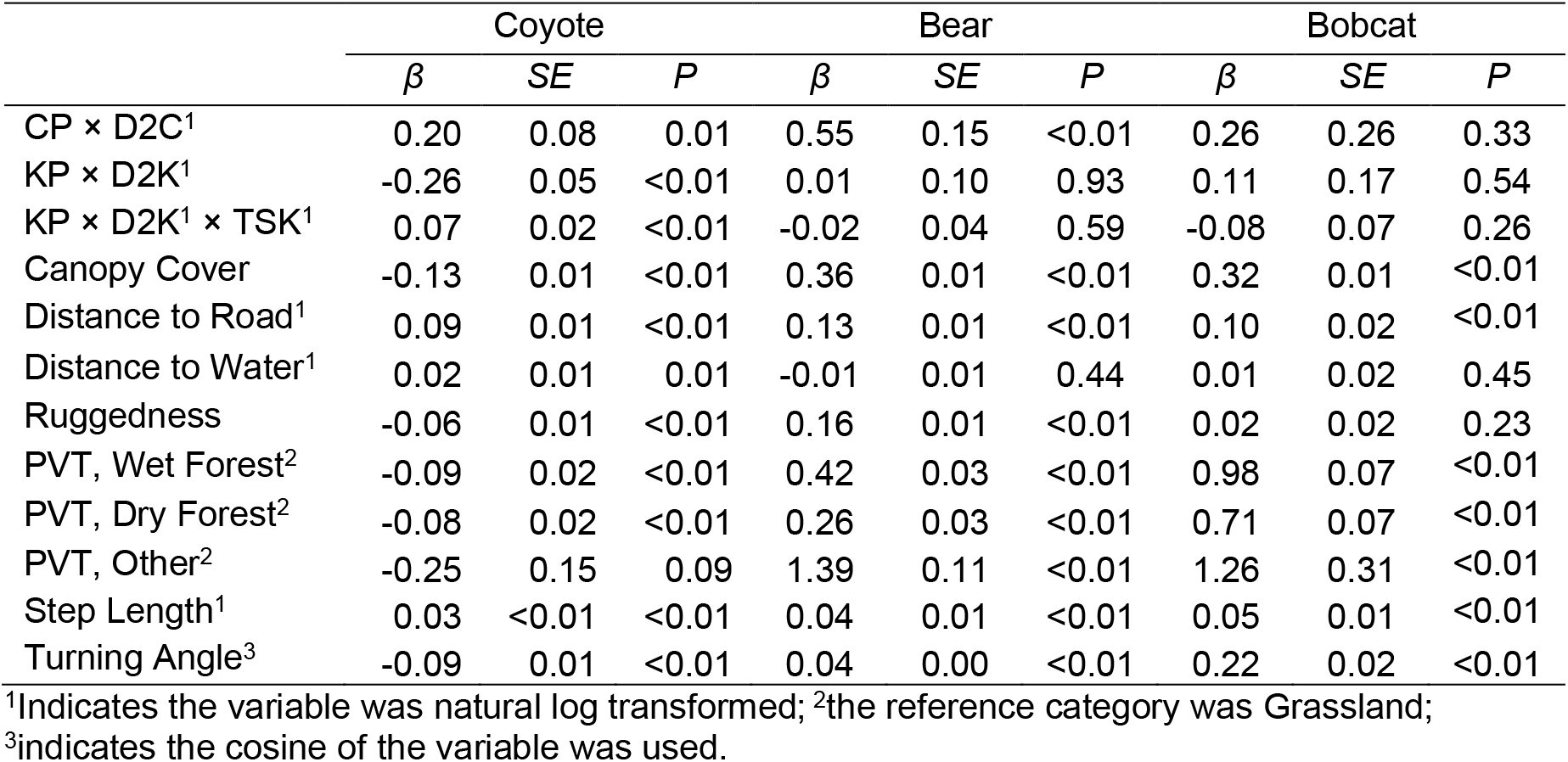
Parameter estimates for landscape and cougar variables influencing the relative probability of selection in step-selection functions in coyotes, black bears, and bobcats. PVT = potential vegetation type, CP = cougar present, D2C = distance to nearest cougar, KP = kill present, D2K = distance to nearest kill, and TSK = time since kill.

**Figure 3.**
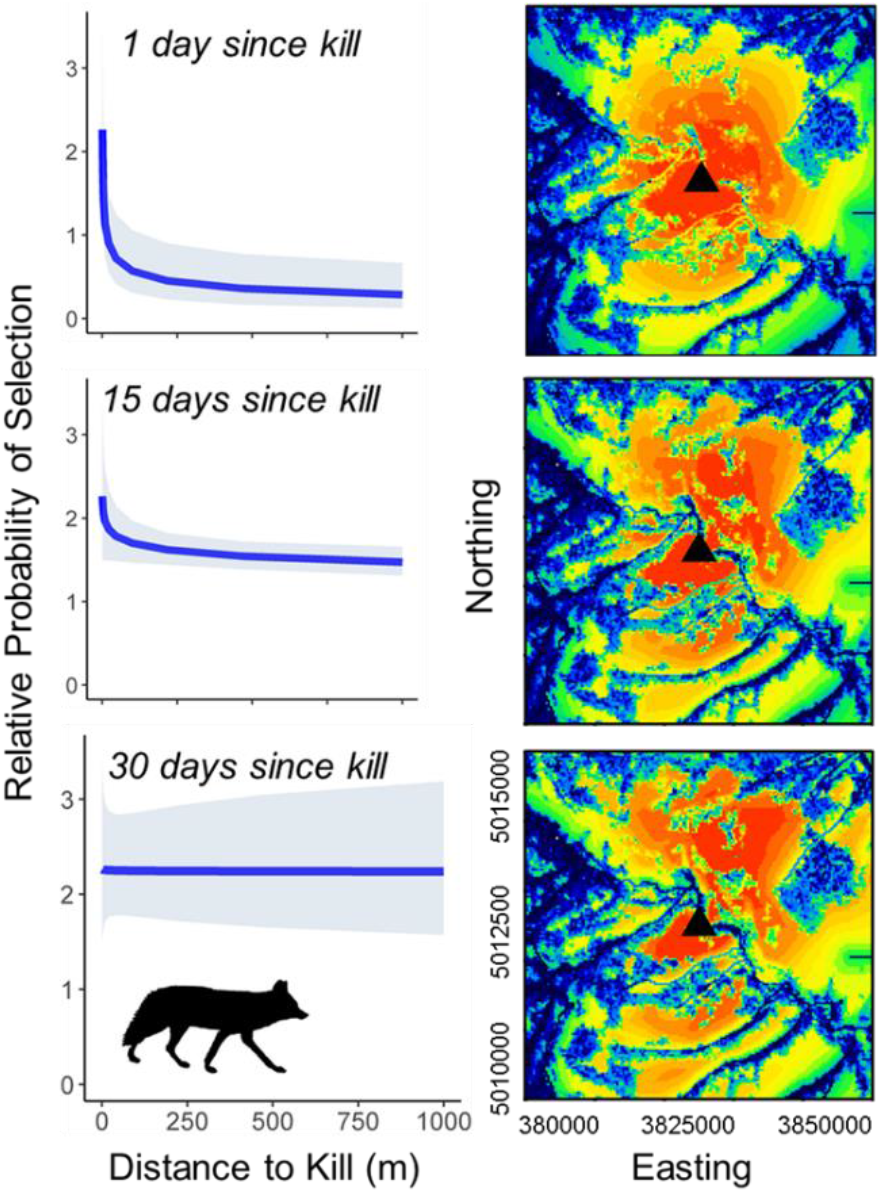
Attraction of coyotes to cougar kill sites. The first column presents marginal response plots for step selection functions showing the relative probability of selection as a function of the distance to the kill site. The second column presents predictive maps showing the relative probability of selecting for areas on the landscape as a function of the distance to the nearest kill site (black triangle). Warm colors indicate selection and cool colors indicate avoidance. In both columns, each row shows the predicted response at 1, 15, and 30 days after the kill was made.

### Diet analysis

DNA metabarcoding of carnivore scats revealed a variable frequency of occurrence of deer and elk across species. In coyotes, elk and deer were present in 58% and 12% of scats, respectively (Fig. 4b). In bears, elk was found in 29% of scats and deer in 8% of scats (Fig. 4b). Zero bobcat scats contained elk and 8% contained deer (Fig. 4b). For cougars, 61% of scats contained elk and 22% contained deer. Investigations of 128 cougar kill sites with confirmed prey revealed that elk represented 64% and deer 16% of the prey items killed by cougars (Figs. 4a, 4b).

**Figure 4.**
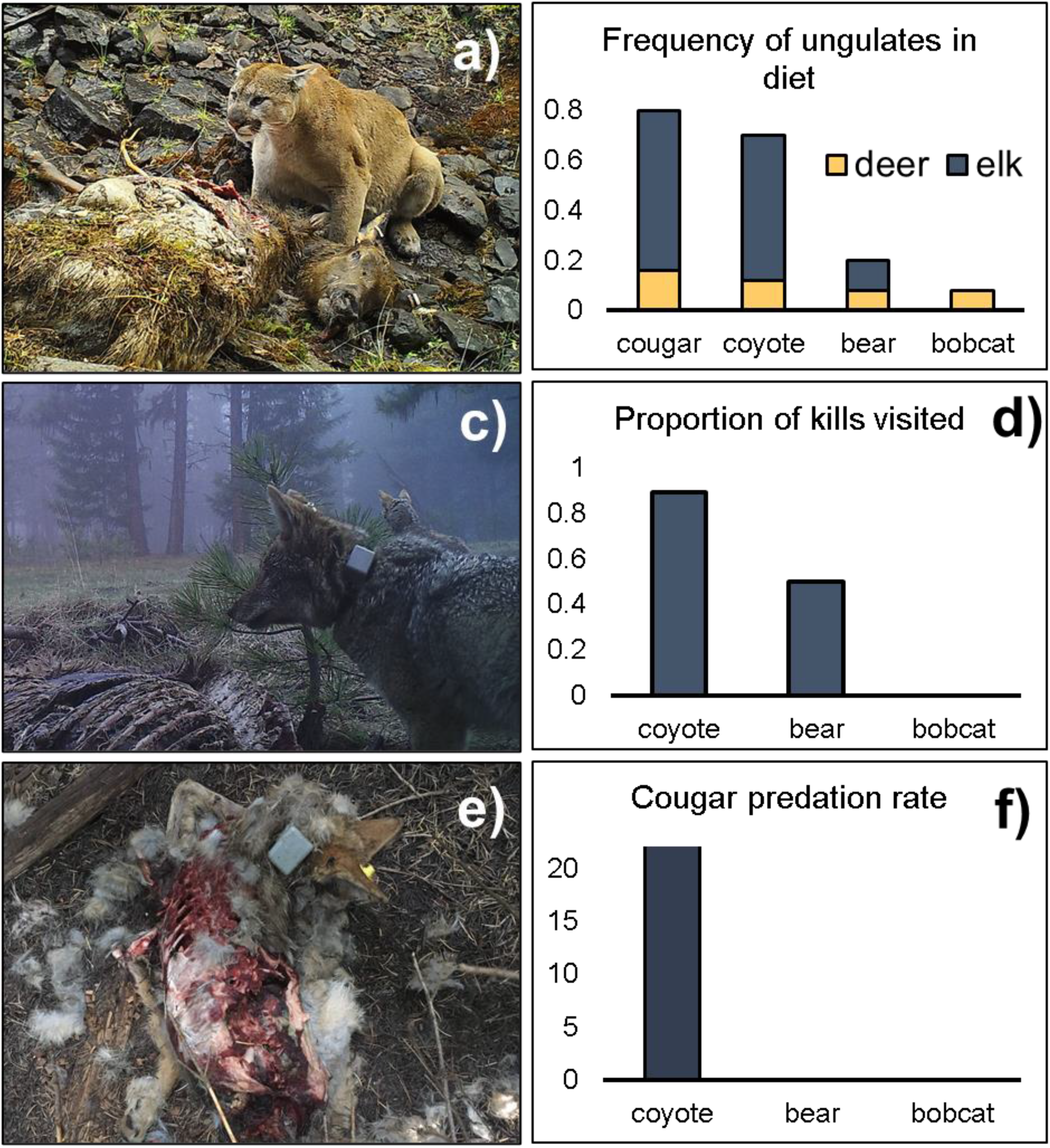
Left panel: Remote camera photos of a) cougar feeding on an adult female elk, c) coyotes scavenging a cougar kill, e): coyote killed by cougar. Right panel: plots showing b) percentages of deer and elk in carnivore diets determined from DNA-metabarcoding of scats (except for cougars in which diet estimates came from kill site investigations), d) proportions of cougar kill sites visited by other carnivores as documented by remote cameras, and f) the estimated proportion of the population of each species killed annually by cougars.

### Scavenging Rates

We monitored 28 cougar kill sites containing ungulate prey using remote cameras to estimate scavenging rates. Cameras were placed an average of 4.4 days after the kill was made (range: 0–16 days; 5 cameras deployed within one day of the kill) and were operational an average of 17.4 (range: 1–52) days. We documented coyotes present at 89% of carcasses, bears at 50% of the carcasses outside the hibernation period (N = 22), and zero bobcats were detected at cougar kills (Fig. 4c, 4d) during the period cameras were active. For coyotes, the negative binomial regression model predicted that one day after a large ungulate was killed, the carcass would receive 1.8 visits by coyotes but that the number of visits would be reduced by 27% with every additional ln(day) (β = −0.31, p = 0.067; Fig. 5a). The coyote visitation rate was expected to be 160% higher for a large ungulate than a juvenile ungulate (β = 0.97, p = 0.059) and was not influenced by cougar presence (β = 0.43, p = 0.26). The bear model predicted 0.36 visits by bears the day after a large ungulate was killed and that every additional ln(day) would reduce the number of visits by 52% (β = −0.73, p = 0.051; Fig. 5b). The number of daily bear visits was not influenced by the size of prey (β = 0.39, p = 0.50) nor whether a cougar was present (β = −0.12, p = 0.89). The intercepts were −0.81 (p = 0.22) and −1.32 (p = 0.29) for the coyote and bear models, respectively.

**Figure 5.**
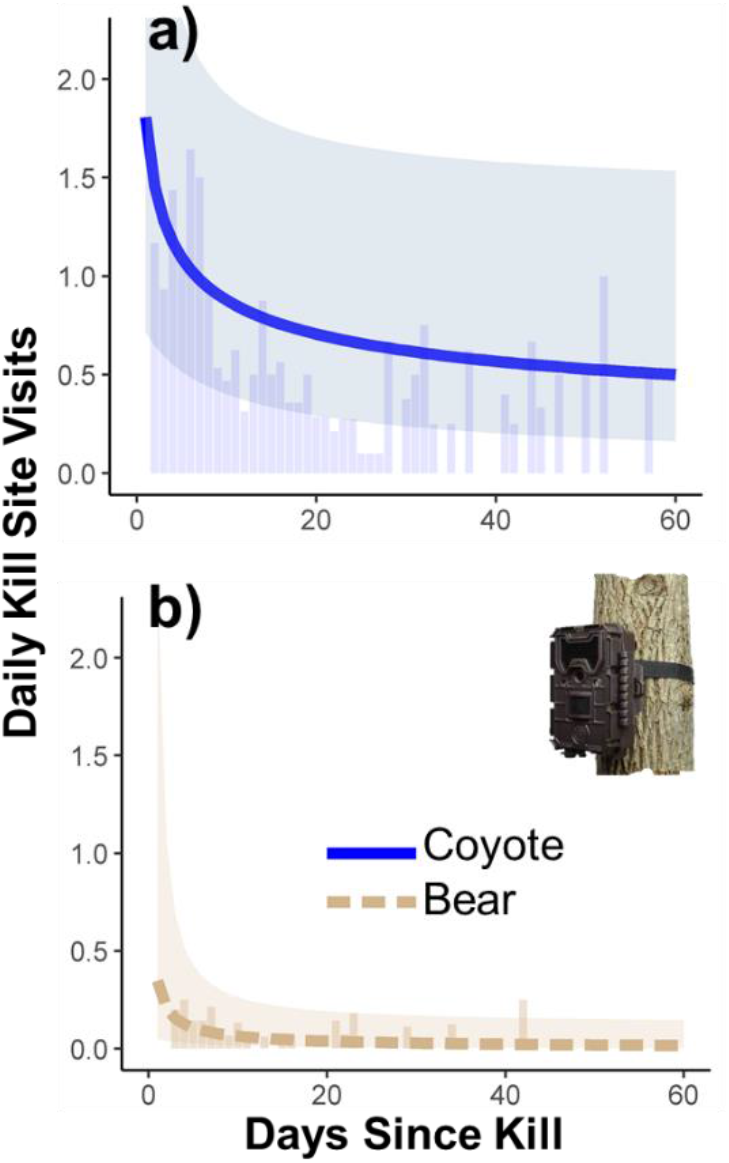
Remote-camera based estimates of daily visits to cougar kill sites by coyotes (panel a) and black bears (panel b). The thick line displays the predicted number of visits for coyotes (solid blue line) and bears (dotted tan line). Shaded regions indicate 95% confidence interval for the predictions. The vertical bars display the mean number of visits for every day since the kill was made, across all kill sites monitored (N = 28). The predictions assume that the kill was a large ungulate (i.e., the binary factor variable for prey size = 1) and that a cougar was present (i.e., the binary factor variable for cougar present = 1).

### Intraguild Predation

Physical investigations of cougar kill sites indicated that coyotes represented 7.0% (9 of 128; 95% CI: 3.3–11.1%) of cougar kills (Fig. 4e, f). In eight cases coyotes were the only prey item found and were consumed by cougars, and in one case a dead coyote was found in conjunction with another prey item. One transient GPS-collared coyote in our sample was killed and consumed by an uncollared cougar outside the study area (Fig. 4e) and in another instance, a GPS-collared cougar killed and consumed a GPS-collared coyote. By observing dead coyotes at 7% of cougar kill sites, and given a cougar density of 2.2 per 100 km^2^ in our study area (22), this suggests that approximately 8.4 coyotes are killed per 100 km^2^ per year. With a coyote density of 33.9 per 100 km^2^ (22), this level of mortality reflects 23.0% (95% CI: 8.4–54.5% when all sources of uncertainty are propagated) of coyotes killed by cougars annually:

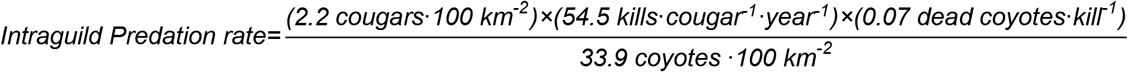

### Weight of Evidence for Facilitation, Suppression, and Null Hypotheses

We found evidence of both facilitation and suppression of other carnivore species by cougars (Fig. 6). Most notably, coyotes exhibited facilitation in three of the five measures evaluated (statistically significant attraction to kill sites, frequent carrion in diet, and high scavenging rates) and indication of suppression in two measures (the presence of intraguild predation and avoidance of cougars). Bears showed evidence of facilitation in two measures (moderate scavenging rates and moderate levels of carrion in diet), suppression in one measure (statistically significant avoidance of cougars), and no evidence in two measures (absence of statistically significant attraction or avoidance of kill sites, absence of intraguild predation). We found no evidence that cougars influenced bobcats in any of the five measures we evaluated, fully supporting the null hypothesis that cougars neither suppress nor facilitate bobcats.

**Figure 6.**
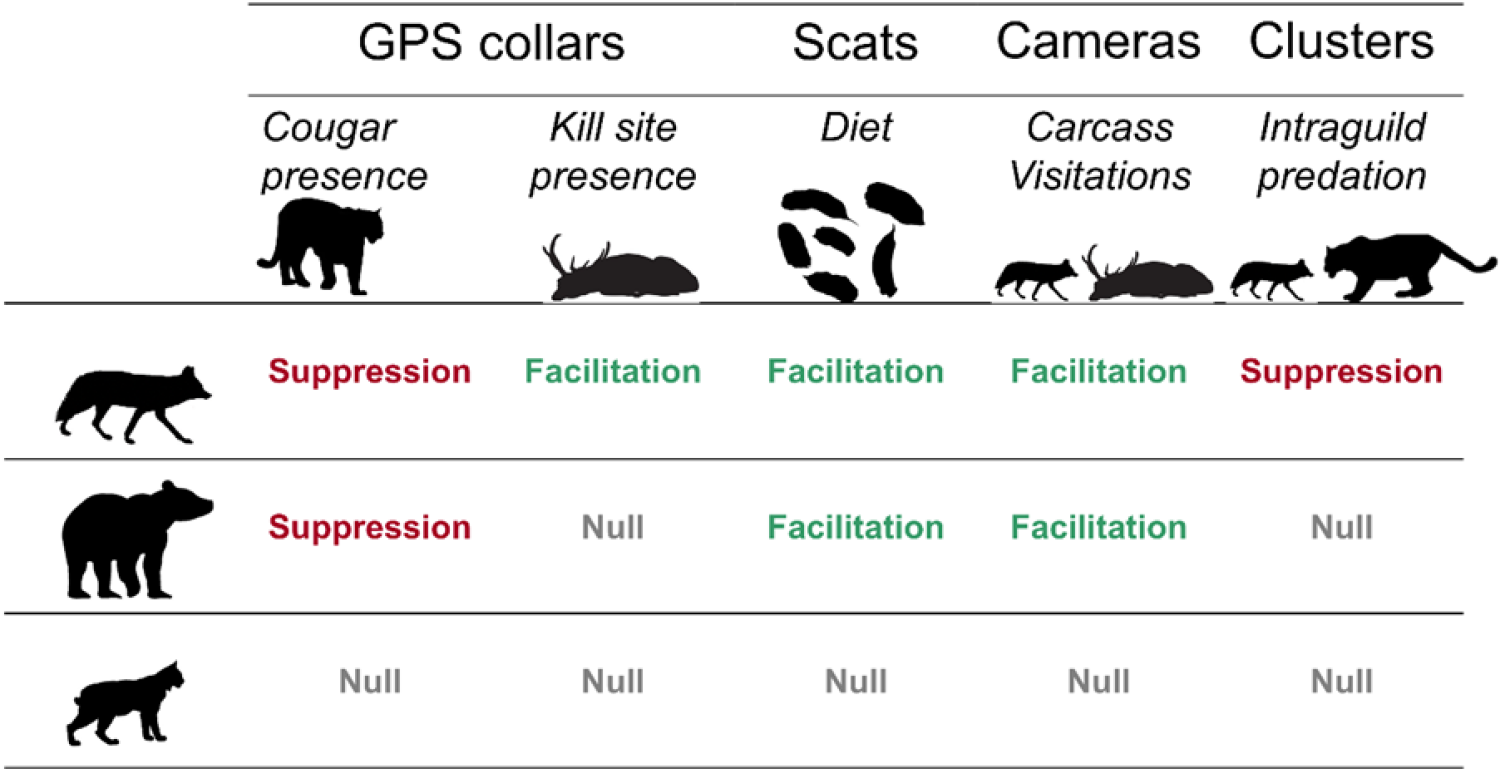
Weight of evidence for support of suppression, facilitation, and null hypotheses regarding the influence of a dominant predator, cougars, on subordinate or co-occurring carnivores: coyotes, bears, and bobcats.

## Discussion

The traditional paradigm concerning species interactions among carnivores has stressed the existence of dominance hierarchies such that dominant predators suppress mesopredators with consequences for their shared prey (1). However, recent work has emphasized the importance of carrion subsidies from dominant to subordinate carnivores. Given the potential for both top-down and bottom-up forcing within the carnivore guild, the generality of whether subordinate species incur a net fitness cost or benefit from dominant predators is far from resolved (7, 12, 14, 37). A particularly missing component has been quantifying both scavenging rate and intraguild predation rate within a given carnivore guild, which a recent meta-analysis found absent from all 256 reviewed studies (7). We assessed the relative strength of suppressive and facilitative forces between a competitively-dominant predator and three subordinate or co-occurring predators by (i) using contemporaneous GPS-tracking of four carnivore species spanning the dominance hierarchy to quantify behavioral avoidance of the dominant species and attraction to its kills, (ii) conducting diet analysis with DNA metabarcoding to quantify potential use of provisioned carrion by subordinate carnivores, (iii) quantifying the visitation rate of subordinate carnivores to the kills of the dominant carnivore, and (iv) by quantifying the strength of top-down interactions measured as the predation rate of dominant carnivores on subordinate species.

Our strongest evidence of facilitation suggested that resource provisioning from cougars to coyotes through scavenging was so ubiquitous as to be an overwhelming determinant of coyote behavior, space use, and resource acquisition. This finding was demonstrated through strong attraction of coyotes to cougar kill sites, coyotes scavenging nearly all carcasses that we monitored, and coyote diets that nearly matched cougars in the magnitude and composition of ungulate consumption. While this clearly suggests that scavenging confers a reward to coyotes, coyotes actively avoided cougars in space, and our estimation that 23% of the coyote population was killed annually by cougars implies that there also exists a strong suppressive effect counteracting the fitness benefit provided by the dominant predator. However, it is not clear that coyotes incurred this predation risk by scavenging given that only 1 of the 9 coyote mortalities caused by cougars occurred in the proximity of a kill site. Nonetheless, this is still suggestive of increased risk while scavenging because coyotes likely spend substantially less than 1/9 of their daily activity budget (2.67 hours daily) at cougar kills. But small sample size precludes statistical determination of how much, or if, predation risk per unit time is higher while actively scavenging, which is needed to determine whether the fatal attraction hypothesis is supported in this system. If risk from dominant predators is not higher at kill sites than elsewhere on the landscape, then scavenging becomes an even more appealing prospect (Fig. 1a). While previous research has found substantial coyote mortality while scavenging at both cougar (21, 38, 39) and wolf kill sites (40), coyotes can also mitigate risk by exhibiting heightened vigilance while scavenging (41) as has also been reported for marten (*Martes martes*) and red fox (*Vulpes vulpes*) at wolf kill sites (42). Two lines of evidence suggest coyotes do not reduce their use of kill sites if cougars are present at the kill; terms accounting for this were not significant in either the regression model of kill site visits from camera data nor a post-hoc step-selection function with an added term indicating whether a cougar was on a kill in question (*SI Appendix*, Text S11, Table S4). This suggests that coyotes readily accept the risk of being near a cougar if a food reward is available, possibly because they can manage the increased risk through vigilance. In addition, if dominant predators become satiated after feeding on a kill, they may have little motivation to kill other carnivores if their primary reason for doing so is to consume them. Determining whether dominant carnivores kill subordinate carnivores for an immediate energetic gain (intraguild predation) or for the long-term benefit of removing a competitor (interference competition) should be the focus of further study.

Theory suggests that different species of carcass competitors should have unique risk-reward calculations even for the same scavenging opportunities (Fig. 1a) and this was clearly true in our system. Not only did GPS tracking indicate no attraction of bears toward cougar kills, scat analysis revealed elk constituted only a moderate percentage of the diets of black bears. Because bears prey on elk neonates (40), an unknown fraction of the elk in their diet may have come from direct predation and not scavenging, leading to an even greater disparity in the amount of carrion consumed between bears and coyotes. A lower rate of bear scavenging at cougar kills is further supported by camera data indicating that bears were less frequent visitors to cougar kills than were coyotes. The lack of attraction to kill sites as measured by GPS data suggests that the bears that did visit cougar-killed prey may have encountered them opportunistically as suggested by previous research (39), in contrast to coyotes which at times made long and directed movements toward kill sites (Fig. S1). While bears neither selected for nor avoided cougar kills, step-selection functions indicated that black bears did actively avoid cougars and this was true for both male and female bears (*SI Appendix*, Text S14, Table S12). This was unexpected because bears are capable of usurping prey from wolves (43), cougars (44), and Eurasian lynx (*Lynx lynx*) (45). However, cougars do kill bears in our study region (21, 37) and literature from other systems reports more instances of cougars killing black bears than the reverse (17). Thus, our results suggesting that black bears viewed cougars as sufficiently threatening such that they made movements to minimize risk of confrontation is defensible. In contrast to coyotes and bears, we found no evidence to suggest that bobcats were in any way influenced by cougars. Their GPS tracks showed no response to cougar presence or kill sites, they never visited a cougar kill monitored by camera, their scats did not contain elk, and they were never found as a prey item at investigations of cougar kill sites. This was surprising given that bobcats scavenge from (15, 39) and fall prey to cougars (38, 46) in other systems, and because empirical evidence suggests that intraguild killing is more likely to occur within taxonomic families (6). By foregoing energetically-profitable cougar kills, and in doing so reducing mortality risk, bobcats in this system are approaching the risk and reward tradeoff quite differently.

So why do subordinate carnivores exhibit such wide variation in behavior toward dominant predators? As illustrated by our dynamic state variable model (Fig. 1a), there are multiple viable solutions to the game of risk and reward. Scavenging only optimizes fitness under a specific set of conditions that depends not only on risk and reward associated with scavenging, but also the probability of finding alternative food sources, the energy required to search for other food, and the current energetic state of the animal. For example, felids expend less energy hunting than do canids (47). If there are ample alternative prey available to bobcats in our system and a low cost of pursuing them, the optimality of eschewing scavenging becomes even more apparent. And of course an animal near starvation should accept more risk to feed on an available carcass than would an animal in prime condition (Fig. 1a), though we lacked the data to assess this. Heightened risk tolerance may also be a function of the timing within an annual cycle. For example, more risk may be accepted to scavenge if increased body condition is needed seasonally to provision young such as is the case for cooperatively breeding canids (e.g. coyotes in Yellowstone National Park had larger litters and higher pup survival by scavenging intensively on elk (40)). Similarly, bears may scavenge more during periods of hyperphagia before hibernation (48). A pattern emerging from other systems is that risk from other scavengers—and not just the owner of the kill—influences the calculus of scavenging given the dominance hierarchy among the carcass competitors. For instance, jackals (*Canis mesomelas* and *Canis aureus*) rarely visit carcasses when there is risk from socially-dominant hyenas (*Crocuta crocuta*) at cheetah (*Acinonyx jubatus*) kills in the Serengeti (49), and hyenas rarely scavenge if male lions (*Panthera leo*) are present (50). Thus, the abstention of bobcats to scavenge in our system may also be due to their position in the dominance hierarchy of carcass competitors since they risk predation from both cougars and numerically-advantaged coyotes, although a post-hoc analysis did not provide support that bobcat made movements to avoid coyotes (*SI Appendix*, Text S12, Table S5). Coyotes in our system faced little risk except from cougars and scavenged profusely; however, recent evidence suggests that their propensity to scavenge is lower in systems where they are subordinate to multiple carcass competitors (37).

The complex interactions among carnivores within a given dominance hierarchy have both direct and indirect effects on species occupying lower trophic levels. There is mounting evidence that scavenging can cause dominant predators to increase their kill rate or modify prey selection. Increased kill rates due to kleptoparasitism have been observed across a wide range of taxa globally, including between eagles (*Haliaeetus leucocephalus*) and falcons (*Falco peregrinus*) in Canada (51), brown bears (*Ursus arctos*) and lynx in central and southeastern Europe (45), and black bears and cougars in North America (15, 16, 44). While we cannot directly assess whether scavenging in our system influenced cougar kill rate, the fact that coyotes routinely scavenged before kill site abandonment by cougars suggests that increased kill rates would be necessary to recover lost biomass (44). In addition to increasing predation rates, there is speculation that scavenging causes predators to select smaller-bodied prey to maximize energetic intake between hunting, feeding, and loss from scavengers, which in turn has evolutionary consequences for carnivore body size (52). Thus, interactions between carnivores can influence prey in complex and indirect ways.

To conclude, it is probably a gross oversimplification to propose that species interactions within terrestrial carnivore communities have either strictly positive, negative, or even neutral consequences (7). We demonstrated that coyotes, and to a lesser extent bears, were confronted by the opposing effects of both facilitation (through food provisioning) and suppression (through death), yet bobcats were seemingly unaffected by the dominant predator. Scavenging carrion from dominant predators incurs both risk and reward, and thus the willingness to engage in this activity should be mediated by behavioral optimization under natural selection. A frontier in carnivore community ecology requires disentangling the net fitness effects of scavenging by subordinate predators. This will be a challenge because mortality itself is an insufficient observation by which to conclude that scavenging incurs a fitness reduction given that it can be optimal to risk death by scavenging (Fig. 1a). Thus, scavenging behavior should be context dependent within and among species given the multiple viable strategies to solve this complex behavioral game subject to the risk and reward tradeoffs unique to each system.

## Acknowledgments

We are grateful to the Oregon Department of Fish and Wildlife, the USDA Forest Service Pacific Northwest Research Station, Oregon State University, and the Wildlife Restoration Act for funding and/or logistical support. We thank a multitude of technicians and volunteers who assisted with fieldwork and photo tagging.

## Notes

**Competing Interest Statement:** The authors declare no competing interest.

### Competing Interest Statement

The authors have declared no competing interest.

### Summary of Updates

The primary modification to this manuscript is a substantial revision of the discussion section based on suggestions arising during peer review.

