## Supporting Information for "Games of risk and reward in carnivore communities"

### **Supplementary Information for**
### Games of risk and reward in carnivore communities

Joel Ruprecht, Charlotte E. Eriksson, Tavis D. Forrester, Derek B. Spitz, Darren A. Clark, Mike J. Wisdom, Marcus Bianco, Mary M. Rowland, Joshua B. Smith, Bruce K. Johnson, and Taal Levi

Joel Ruprecht


**This PDF file includes:**

Supplementary text
Figures S1 to S2
Tables S1 to S11
Legends for Movies S1 to S3
SI References

**Other supplementary materials for this manuscript include the following:**

Movies S1 to S3

**Supplementary Information Text**

**Text S1. Development of a dynamic state variable model investigating risk and reward of scavenging.** We developed a two-patch dynamic state variable model (1) to qualitatively understand the conditions under which scavenging versus hunting alternative prey could represent the optimal decision for a carnivore seeking to maximize fitness. Specifically, we investigated whether the reward accrued from scavenging (i.e., facilitation) could optimize fitness despite a heightened level of risk from intraguild predation (suppression). When carnivores are given a chance to scavenge from a dominant species or competitor, they can either disregard the opportunity and continue to hunt alternative prey (representing the choice of Patch 1) or choose to scavenge carrion from the kill, readily gaining access to food but perhaps incurring a higher mortality risk (Patch 2). Under state-dependent life history theory, the choice of patch *i* represents the option that maximizes fitness (*F*) and depends on the animal's energetic state (*x*) at a given week (*t*) within a biological year, weekly energetic costs (c), the probability of finding food in patch *i* ($\lambda_i$), the probability of dying in patch *i* ($d_i$, i.e., *risk,*), and the profitability of patch *i* ($p_i$, i.e. *reward*). The fitness value for visiting patch *i* ($V_i$) is given by:

$$V_i = (1-d_i)\big(\lambda_i F(x-c_i+p_i,\ t+1) + (1-\lambda_i)F(x-c_i,\ t+1)\big)$$

We assumed that fitness was a saturating function of energetic state such that:

$$F_t = \frac{x_t-c}{x_t-c+b}$$

where *b* is the energetic state in which fitness is half its maximum and $x_t - c$ is the energetic state at week *t* after energetic expenditures. We let energetic state range from 1–100 where values ≤ 10 indicated death from starvation. We varied the probability of finding food from hunting and the risk incurred from scavenging to illustrate how different strategies could become optimal under different scenarios. We used values of *d* (daily probability of death) of 0.002 for Patch 1 (hunting alternative prey) and 0.002, 0.005, and 0.01 (imputing annual survival rates of 0.9, 0.75, and 0.6, respectively) for Patch 2 (scavenging). We varied the probability of finding food ($\lambda$), from 0.4 to 0.8 for Patch 1 and held this value constant at 0.9 for Patch 2. For both patches, we assumed that daily energetic costs (*c*) were 2 units and that profitability (*p*) was 4 units if food was found.

The stochastic dynamic programming equation (1) identifies the optimal choice of patch (i.e. scavenging or hunting alternative prey) at a given week *t* in an annual cycle and for a given energetic state *x*:

$$F(x,t) = \max(V_1(x,t),\ V_2(x,t))$$

Where $V_1$ is the value of hunting alternative prey and $V_2$ is the value of scavenging.

**Text S2. Methods for capture and GPS collaring of carnivores.** We captured and GPS-collared each of the four carnivore species as part of concurrent research on predator-prey relationships. Coyotes were captured using padded foothold traps immobilized with tiletamine-zolazepam (Telazol®) at a concentration of 10 mg/kg. A GPS collar (Lotek MiniTrack, Lotek Wireless Inc., Newmarket, ON, Canada or Vectronic Vertex, Vectronic Aerospace GmbH, Berlin, Germany) was placed on each adult coyote and was programmed to record locations every 2 or 3 hours. We captured bobcats using cage traps baited with visual and olfactory attractants and administered ketamine (10 mg/kg) and xylazine (1.5 mg/kg) for immobilization, and upon release, yohimbine (0.125 mg/kg; Yobine®) was given as an antagonist for xylazine. Each bobcat was fit with a GPS collar (Lotek MiniTrack, Lotek Wireless Inc., Newmarket, ON, Canada) scheduled to take fixes every 2 hours. Black bears were captured using culvert traps or padded foot snares and were immobilized with Telazol® at a concentration of 7 mg/kg. Bears were fit with GPS collars (Lotek GPS 7000 or LiteTrack Iridium 420, Lotek Wireless Inc., Newmarket, ON, Canada) that recorded positions every 2 hours. We captured cougars using trained pursuit hounds. We

searched for fresh cougar tracks in snow along roads within the study area and when located, released hounds to pursue tracks until the cougar was treed. When treed, cougars were immobilized with ketamine (10 mg/kg) and xylazine (2 mg/kg) via dart gun, and before release administered yohimbine (0.125 mg/kg; Yobine®) as an antagonist for xylazine. A GPS collar (Lotek GPS 4400S, IridiumTrack M, Lotek Wireless Inc., Newmarket, ON, Canada or Vectronic Vertex Lite, Vectronic Aerospace GmbH, Berlin, Germany) was placed on each cougar and was programmed to record locations every 3 hours. For all species, GPS data were screened and errant locations were removed using established protocol for cleaning GPS data (2).
All animal handling was performed in accordance with protocols approved by the USDA Forest Service, Starkey Experimental Forest Institutional Animal Care and Use Committee (IACUC No. 92-F-0004; protocol #STKY-16-01) and followed the guidelines of the American Society of Mammalogists for the use of wild mammals in research (3).

**Text S3. Study area description.** Our study was centered at Starkey Experimental Forest and Range in the Blue Mountains of northeastern Oregon (45.247, -118.563). The study area is composed of a patchwork of ponderosa pine (*Pinus ponderosa*) stands and mixed pine-fir forests (*Pinus*, *Abies* and *Pseudotsuga* spp.), punctuated with grasslands dominated by native bunchgrasses (*Poa*, *Danthonia*, and *Pseudoroegneria* spp.) and invasive annual grasses (*Bromus* and *Ventenata* spp.) (4). Elevation ranges between 1,122 and 1,500 m in the study area, where an average of 51 cm of precipitation falls annually, typically as snow in the winter months (4).

**Text S4. Detection dog surveys for scat collection.** Scat detection dogs from the University of Washington Conservation Canine program surveyed a 224 $km^2$ study area between 6 and 26 June 2017. The area surveyed was composed of 56 grid cells each with an area of 4 $km^2$. Detection dogs surveyed 6–8 km linear distance within each cell to distribute effort across the study area. Dog handlers were not given specific survey routes but were encouraged to follow natural travel corridors such as ridgelines, saddles, drainage bottoms, game trails, and fence lines. No more than 50% of the distance traveled per gridcell was permitted to be on linear features such as trails or roads. When scats were located, the handler recorded the GPS position and placed the entire scat in triplicate paper bags. Within 72 hours of collection, scats were desiccated in a drying oven for 24 hours at 40°C (5). Detection dogs in our study were trained to locate black bear, coyote, cougar, and bobcat scats.

**Text S5. Methods of determining species ID and DNA metabarcoding of scats for diet analysis**. We used DNA metabarcoding to characterize the presence of vertebrate prey items in the scats of bears, bobcats, and coyotes. We extracted DNA from the scats (15 samples per extraction batch) using the DNeasy Blood and Tissue kit (Qiagen, USA) and included 1 extraction blank per batch. The prey and defecator of each scat was identified by amplifying ~100 bp of the mitochondrial 12S gene region (primers used in (6), (7), and modified from (8)). Each PCR reaction was amplified with identical unique 8 bp tags on the 5' end of the forward and reverse primers to allow for sample identification and to prevent tag-jumping (9). We performed 3 PCR replicates per scat and monitored for contamination by including 3 no-template controls per 96-well plate. PCR was performed in 20 μL reactions using 10 μL Kapa HiFi HotStart High Fidelity ReadyMix (Kapa Biosystems), 5.6 μL of forward and reverse primers (0.25 μM final concentration), 2.4 μL of water, and 2 μL DNA extracts (including extraction blanks and PCR no template controls). Cycling conditions were 95˚C initial denaturation for 3 minutes, followed by 35 cycles of 98˚C for 20 seconds, 58˚C for 15 seconds, 72˚C for 30 seconds, and a final extension at 72˚C for 1 minutes. Following PCR, we quantified the DNA concentration of each sample using a fluorescence microplate reader with the AccuBlue dsDNA Quantitation Kit (Biotium, USA) and normalized each sample accordingly. We then pooled 3 μL from each sample into a 0.65 mL Eppendorf tube per 96-well plate. We used NEBNext Ultra II Library Prep Kit (New England Biolabs, USA) to adapt the library pools into Illumina sequencing libraries following the manufacturer's instructions (Illumina Inc, USA). We then purified the library pools using PCRClean DX (Aline Biosciences, USA), quantified DNA concentration using a Qubit 2.0

fluorometer (Life Technologies, USA), and normalized each pool before sending the libraries for 150 bp paired-end sequencing on an Illumina HiSeq 3000 at the Center for Genome Research and Biocomputing, Oregon State University. Library size distribution was checked prior to sequencing using a High Sensitivity D5000 DNA ScreenTape assay on an Agilent Tapestation 4200 (Agilent Technologies, USA). The raw sequence reads were paired using PEAR (10) and demultiplexed based on the unique 8 bp index tags using a custom shell script. Unique reads from each sample replicate were clustered, counted and taxonomically assigned using BLAST (www.ncbi.nlm.nih. gov/blast), against all 12S vertebrate sequences in Genbank. We used the negative controls to set filtering read thresholds and assigned species if present in at least 2 out of the 3 replicates.

**Text S6. Identification of cougar kill sites.** After killing large prey, cougars remain close to the carcass for several days as they feed (11). This reduction in movement can be detected in GPS collar locations, which often display a distinct cluster of locations that can be distinguished from other behaviors such as traveling or searching for prey (12, 13). We physically investigated a subset of potential cougar kills, hereafter “clusters” and confirmed prey at 128 sites following the protocol outlined by (13), and the remainder were identified using a clustering algorithm developed by (12) and modified by (13). However, that algorithm was developed to identify potential (not probable) kill sites and therefore generated an unacceptably high number of false positives to meet our objectives. We therefore used logistic regression equations developed by (13) and (14) to predict the probability of the presence of an ungulate prey item at the cluster (1/0) to restrict potential clusters to those with a high probability of containing ungulate prey. We validated the model by comparing its predictions with clusters physically investigated and determined that the model correctly identified true positives (i.e. the presence of an ungulate prey item) in 81.6% of cases and identified true negatives in 86.5% of instances.

**Text S7. Details on predation rate calculations.** We calculated the cougar intraguild predation rate on coyotes ($P_{coyote}$) as the number of coyotes killed per unit time ($K_{coyote}$) divided by coyote population density ($D_{coyote}$), where $K_{coyote}$ is the product of cougar population density ($D_{cougar}$), cougar kill rate (kills/year; $R$), and proportion of kills corresponding to coyotes ($F_{coyote}$):

$$P_{coyote} = \frac{K_{coyote}}{D_{coyote}} = \frac{D_{cougar} \times R \times F_{coyote}}{D_{coyote}}$$

Each parameter in the predation rate equation was estimated using empirical data and therefore is itself a random variable with an associated measure of uncertainty that we sought to acknowledge. Thus, in order to propagate all the error in predation rate calculation arising from the uncertainty of each individual parameter, we used a Monte Carlo simulation approach. We calculated the predation rate using 10,000 Markov Chain Monte Carlo simulations where each iteration used a random draw from a distribution matching the mean and variance of the empirical estimates for each parameter. We used gamma distributions for cougar and coyote densities from a genetic spatial-capture recapture study in the same area (15), a gamma distribution for number of cougar kills per year from an adjacent study area (13), and a beta distribution for the proportion of cougar kills in which coyotes were the prey based on kill site investigations in this study:

$$\text{cougar density} \sim \text{Gamma}\left(\frac{2.2^2}{0.7^2}, \frac{2.2}{0.7^2}\right)$$

$$\text{coyote density} \sim \text{Gamma}\left(\frac{33.9^2}{3.3^2}, \frac{33.9}{3.3^2}\right)$$

$$\text{kill rate} \sim \text{Gamma}\left(\frac{54.5^2}{3.0^2}, \frac{54.5}{3.0^2}\right)$$

$$\text{proportion coyote} \sim \text{Beta}(9, 119)$$

The distributions specified above assume 1.7 (standard deviation = 0.7) cougars per 100 $km^2$, 25.0 (standard deviation = 2.6) coyotes per 100 $km^2$, a cougar kill rate of 54.5 (standard deviation = 3.0) prey items per year, and that coyotes represented 7.0% of the prey items killed by cougars (9 coyotes out of 128 prey items).

**Text S8. Description of spatial covariates used in SSFs.** We calculated the natural log-transformed Euclidean distances between the endpoint of each observed and random step and the nearest road open to motorized vehicle use and perennial water source (data sources from the US Forest Service). We used Integrated Land Assessment Project products for estimates of canopy cover (16) and delineations of potential vegetation type (17), a factor variable with categories for wet forest, dry forest, grasslands, and other. We used the vector ruggedness measure (18) to characterize variation in terrain, aspect, and slope. All continuous spatial variables were scaled to have a mean of zero and standard deviation of one.

**Text S9. Evaluation of coyote step-selection functions separated by sex.** Post-hoc, we evaluated whether attraction to known or predicted cougar kill sites varied as a function of sex in coyotes by fitting SSFs separately for males ($N$ = 10) and females ($N$ = 7). The direction of the D2C and D2K variables were similar for both males and females, although the magnitude of these terms were smaller and not statistically significant in the female-only model (Table S2).

**Text S10. Evaluation of coyote step-selection functions separated by resident vs transient status.** Post-hoc, we evaluated whether attraction to known or predicted cougar kill sites varied as a function of the behavioral status (i.e. resident vs transient) in coyotes. Coyotes were classified as residents ($N$ = 10) if they were faithful to small (< 20 $km^2$) territories and exhibited central-point foraging behavior, or transient ($N$ = 7) if they had large (200–400 $km^2$) or poorly-defined territories and exhibited random, transitory movements without fidelity to any given area. The direction and magnitude of the D2C and D2K variables were similar between residents and transients, although these terms were not statistically significant in the transient-only model (Table S3).

**Text S11. Evaluation of a coyote step-selection function with a term for 'cougar on kill'.** Post-hoc, we evaluated whether the inclusion of a variable indicating whether the nearest cougar was on a kill (hereafter, 'cougar on kill' or CK) modified the effect of D2K. CK was a binary indicator variable denoting whether the nearest cougar was within 100 m of a known or predicted kill. We cast CK as an interaction with D2K allowing the sign and magnitude of D2K to differ based on whether the cougar was on a kill. In this model, D2K as a main effect remained significant ($p$ = < 0.001) indicating coyotes are attracted to kill sites (Table S4). The interaction between CK and D2K was not significant ($p$ = 0.81) suggesting that coyotes were no less likely to visit the kill if a cougar was present (Table S4). We suggest this provides evidence that coyotes avoid cougars when they offer no food reward but accept the risk of being near a cougar if scavenging opportunities are available.

**Text S12. Evaluation of a step-selection function assessing bobcat avoidance of coyotes.** Post-hoc, we evaluated whether bobcats altered movements in response to the nearest known coyote (Table S5). We fit SSFs for bobcats with the same suite of landscape and movement variables as the other analyses but we replaced "Distance to Cougar" with "Distance to Coyote" (D2C). The D2C variable measured whether bobcats made movements toward versus away from the nearest known coyote if one was present within 1,000 m of the focal bobcat ("Coyote Present" or CP). This analysis was motivated by our finding that bobcats were not attracted to cougar kill sites and we thereby sought to determine whether coyote presence could explain the lack of attraction to kill sites since coyote activity dominated scavenging activity at carcasses.

**Text S13. Evaluation of different buffer sizes to determine "cougar presence" or "kill presence" in step-selection functions.** To ensure that inference on the direction or magnitude of the effects of D2C and D2K in step selection functions was not influenced by the choice of buffers indicating whether a cougar or kill was present, we evaluated other potential buffers. For cougar presence (CP) our original analysis used a value of 1,000 m beyond which we assumed a cougar could not be detected by a co-occurring carnivore. We also present results for buffers of 500 m and 1,500 m (Table S6–S8). Similarly, for kill presence (KP) we evaluated values of 2,000 m and 4,000 m (Table S9–S11) in addition to our original analysis using 3,000 m.

**Text S14. Evaluation of black bear step-selection functions separated by sex.** Post-hoc, we evaluated whether attraction to known or predicted cougar kill sites varied as a function of sex in bears by fitting SSFs separately for males ($N$ = 7) and females ($N$ = 4). While the D2K variable was not significant for either males or females, interpreting the sign of the coefficients suggested that males tended to be attracted to kill sites whereas females tended to avoid them (Table S12).

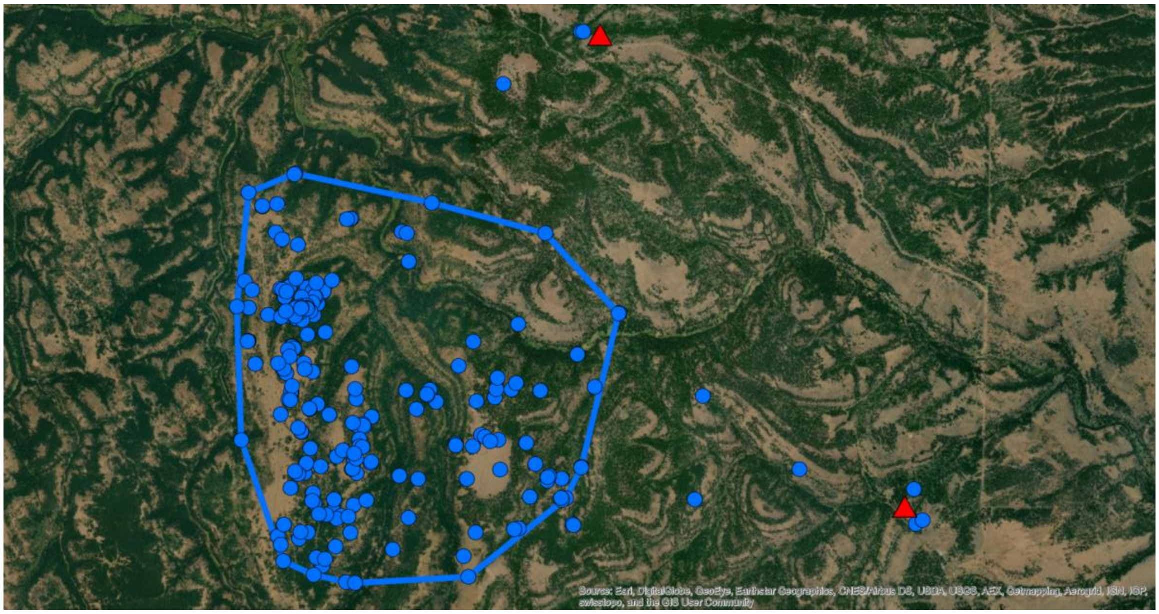


**Fig. S1.** GPS locations showing two separate bouts of male coyote L13 traveling to reach an elk carcass (red triangle). The movements were 2.2 km and 3.6 km from the nearest perimeter of the animal’s home range as defined by a 95% minimum convex polygon (blue polygon). Blue points indicate GPS collar locations.

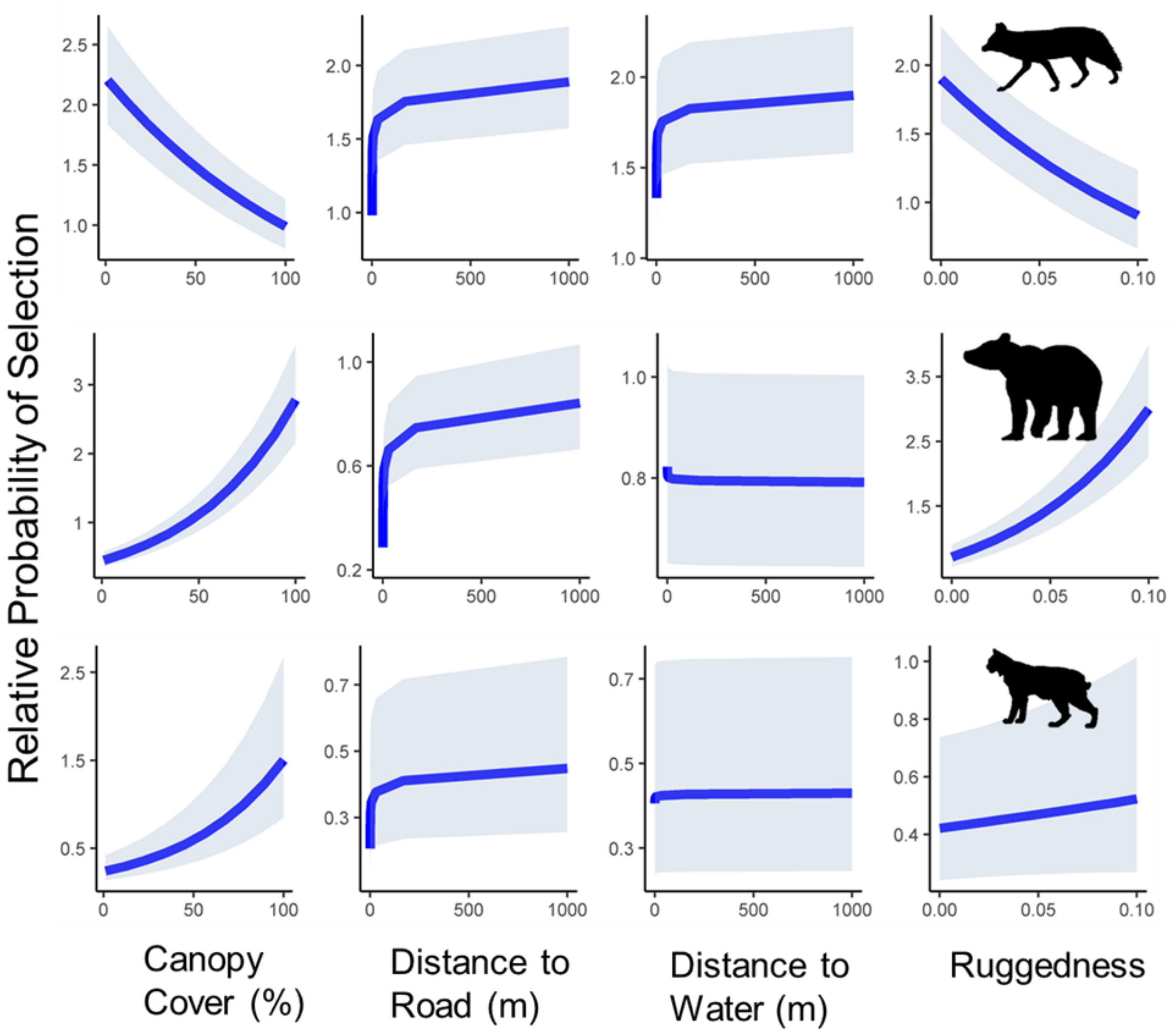


**Fig. S2.** Marginal response plots showing the relative probability of selection of landscape features estimated from step selection functions for (top to bottom) coyotes, black bears, and bobcats. The model estimate is displayed by the blue line and the 95% confidence interval is provided by the shaded band.

**Table S1.** Model selection results of step-selection functions for habitat-only models and habitat + cougar models. ΔAIC gives the change in AIC value from the top model where values of zero indicate the top model.

| Species | ΔAIC | |
|---|---|---|
| | Habitat Only | Habitat + Cougar |
| Bear | 10.36 | 0 |
| Bobcat | 0 | 2.69 |
| Coyote | 24.0 | 0 |

**Table S2.** Parameter estimates for step-selection functions for coyotes separated by sex. PVT = potential vegetation type, CP = cougar present, D2C = distance to nearest cougar, KP = kill present, D2K = distance to kill, and TSK = time since kill.

| | Male | | | Female | | |
|---|---|---|---|---|---|---|
| | *β* | *SE* | *P* | *β* | *SE* | *P* |
| Canopy Cover | -0.14 | 0.010 | <0.001 | -0.09 | 0.02 | <0.001 |
| Distance to Road[1] | 0.087 | 0.011 | <0.001 | 0.08 | 0.02 | <0.001 |
| Distance to Water[1] | 0.026 | 0.011 | 0.017 | 0.01 | 0.02 | 0.44 |
| Ruggedness | -0.059 | 0.011 | <0.001 | -0.08 | 0.02 | <0.001 |
| PVT, Wet Forest[2] | -0.037 | 0.029 | 0.20 | -0.18 | 0.04 | <0.001 |
| PVT, Dry Forest[2] | -0.066 | 0.024 | 0.0065 | -0.09 | 0.04 | 0.03 |
| PVT, Other[2] | -0.19 | 0.19 | 0.31 | -0.35 | 0.25 | 0.16 |
| Step Length[1] | 0.028 | 0.0038 | <0.001 | 0.02 | 0.01 | <0.001 |
| Turning Angle[3] | -0.11 | 0.011 | <0.001 | -0.05 | 0.02 | 0.01 |
| CP × D2C[1] | 0.25 | 0.10 | 0.012 | 0.10 | 0.15 | 0.52 |
| KP × D2K[1] | -0.31 | 0.063 | <0.001 | -0.13 | 0.11 | 0.25 |
| KP × D2K[1] × TSK[1] | 0.086 | 0.027 | 0.0012 | 0.03 | 0.05 | 0.52 |

[1]Indicates the variable was natural log transformed; [2]the reference category was Grassland; [3]indicates the cosine of the variable was used.

**Table S3.** Parameter estimates for step-selection functions for coyotes separated by behavioral status (resident vs transient). PVT = potential vegetation type, CP = cougar present, D2C = distance to nearest cougar, KP = kill present, D2K = distance to kill, and TSK = time since kill.

| | Resident | | | Transient | | |
|---|---|---|---|---|---|---|
| | *β* | *SE* | *P* | *β* | *SE* | *P* |
| Canopy Cover | -0.13 | 0.01 | <0.001 | -0.12 | 0.02 | <0.001 |
| Distance to Road[1] | 0.08 | 0.01 | <0.001 | 0.09 | 0.02 | <0.001 |
| Distance to Water[1] | 0.04 | 0.01 | <0.001 | -0.01 | 0.02 | 0.53 |
| Ruggedness | -0.06 | 0.01 | <0.001 | -0.06 | 0.02 | <0.001 |
| PVT, Wet Forest[2] | -0.07 | 0.03 | 0.01 | -0.11 | 0.04 | 0.01 |
| PVT, Dry Forest[2] | -0.08 | 0.03 | <0.001 | -0.07 | 0.04 | 0.06 |
| PVT, Other[2] | -0.29 | 0.25 | 0.24 | -0.25 | 0.19 | 0.18 |
| Step Length[1] | 0.03 | <0.001 | <0.001 | 0.01 | 0.01 | 0.03 |
| Turning Angle[3] | -0.16 | 0.01 | <0.001 | 0.04 | 0.02 | 0.02 |
| CP × D2C[1] | 0.22 | 0.10 | 0.02 | 0.17 | 0.16 | 0.29 |
| KP × D2K[1] | -0.30 | 0.06 | <0.001 | -0.10 | 0.13 | 0.47 |
| KP × D2K[1] × TSK[1] | 0.09 | 0.03 | <0.001 | -0.01 | 0.06 | 0.90 |

[1]Indicates the variable was natural log transformed; [2]the reference category was Grassland; [3]indicates the cosine of the variable was used.

**Table S4.** Parameter estimates for step-selection functions for coyotes with an added term for ‘cougar on kill’ (CK) that evaluates whether the sign or magnitude of ‘distance to kill’ changes if a cougar is 100 m from the kill. This model was constructed to assess whether coyotes are less attracted to cougar kill sites if the cougar is present. PVT = potential vegetation type, CP = cougar present, D2C = distance to nearest cougar, KP = kill present, D2K = distance to kill, and TSK = time since kill.

| | *β* | *SE* | *P* |
|---|---|---|---|
| Canopy Cover | -0.13 | 0.01 | <0.001 |
| Distance to Road[1] | 0.09 | 0.01 | <0.001 |
| Distance to Water[1] | 0.02 | 0.01 | 0.01 |
| Ruggedness | -0.06 | 0.01 | <0.001 |
| PVT, Wet Forest[2] | -0.09 | 0.02 | <0.001 |
| PVT, Dry Forest[2] | -0.08 | 0.02 | <0.001 |
| PVT, Other[2] | -0.25 | 0.15 | 0.09 |
| Step Length[1] | 0.03 | <0.001 | <0.001 |
| Turning Angle[3] | -0.09 | 0.01 | <0.001 |
| CP × D2C[1] | 0.21 | 0.08 | 0.01 |
| KP × D2K[1] | -0.26 | 0.06 | <0.001 |
| KP × D2K[1] × TSK[1] | 0.07 | 0.02 | <0.001 |
| CK × D2K[1] | -0.01 | 0.05 | 0.81 |

[1]Indicates the variable was natural log transformed; [2]the reference category was Grassland; [3]indicates the cosine of the variable was used.

**Table S5.** Parameter estimates for step-selection functions for bobcats with covariates for landscape variables and the presence of the nearest known coyote. This model was constructed to assess whether bobcats alter movements in response to coyote presence. PVT = potential vegetation type, CP = coyote present, and D2C = distance to nearest known coyote.

| | $\beta$ | *SE* | *P* |
|---|---|---|---|
| Canopy Cover | 0.32 | 0.01 | <0.001 |
| Distance to Road[1] | 0.10 | 0.02 | <0.001 |
| Distance to Water[1] | 0.01 | 0.02 | 0.58 |
| Ruggedness | 0.01 | 0.02 | 0.42 |
| PVT, Wet Forest[2] | 0.95 | 0.07 | <0.001 |
| PVT, Dry Forest[2] | 0.67 | 0.07 | <0.001 |
| PVT, Other[2] | 1.38 | 0.32 | <0.001 |
| Step Length[1] | 0.05 | 0.01 | <0.001 |
| Turning Angle[3] | 0.21 | 0.02 | <0.001 |
| CP × D2C[1] | 0.06 | 0.05 | 0.25 |

[1]Indicates the variable was natural log transformed; [2]the reference category was Grassland; [3]indicates the cosine of the variable was used.

**Table S6.** Parameter estimates for step selection functions for coyotes using different buffer sizes to determine cougar presence (CP). PVT = potential vegetation type, D2C = distance to nearest cougar, KP = kill present, D2K = distance to kill, and TSK = time since kill.

| | 500 m | | | 1500 m | | |
|---|---|---|---|---|---|---|
| | *β* | *SE* | *P* | *β* | *SE* | *P* |
| Canopy Cover | -0.13 | 0.01 | <0.001 | -0.13 | 0.01 | <0.001 |
| Distance to Road[1] | 0.09 | 0.01 | <0.001 | 0.09 | 0.01 | <0.001 |
| Distance to Water[1] | 0.02 | 0.01 | 0.01 | 0.02 | 0.01 | 0.01 |
| Ruggedness | -0.06 | 0.01 | <0.001 | -0.06 | 0.01 | <0.001 |
| PVT, Wet Forest[2] | -0.09 | 0.02 | <0.001 | -0.09 | 0.02 | <0.001 |
| PVT, Dry Forest[2] | -0.08 | 0.02 | <0.001 | -0.08 | 0.02 | <0.001 |
| PVT, Other[2] | -0.25 | 0.15 | 0.09 | -0.25 | 0.15 | 0.09 |
| Step Length[1] | 0.03 | <0.001 | <0.001 | 0.03 | <0.001 | <0.001 |
| Turning Angle[3] | -0.09 | 0.01 | <0.001 | -0.09 | 0.01 | <0.001 |
| CP × D2C[1] | 0.35 | 0.12 | <0.001 | <0.001 | 0.05 | 0.97 |
| KP × D2K[1] | -0.26 | 0.05 | <0.001 | -0.22 | 0.06 | <0.001 |
| KP × D2K[1] × TSK[1] | 0.07 | 0.02 | <0.001 | 0.06 | 0.02 | 0.02 |

[1]Indicates the variable was natural log transformed; [2]the reference category was Grassland; [3]indicates the cosine of the variable was used.

**Table S7.** Parameter estimates for step selection functions for black bears using different buffer sizes to determine cougar presence (CP). PVT = potential vegetation type, D2C = distance to nearest cougar, KP = kill present, D2K = distance to kill, and TSK = time since kill.

| | 500 m | | | 1500 m | | |
|---|---|---|---|---|---|---|
| | *β* | *SE* | *P* | *β* | *SE* | *P* |
| Canopy Cover | 0.36 | 0.01 | <0.001 | 0.36 | 0.01 | <0.001 |
| Distance to Road[1] | 0.14 | 0.01 | <0.001 | 0.14 | 0.01 | <0.001 |
| Distance to Water[1] | -0.01 | 0.01 | 0.37 | -0.01 | 0.01 | 0.37 |
| Ruggedness | 0.16 | 0.01 | <0.001 | 0.16 | 0.01 | <0.001 |
| PVT, Wet Forest[2] | 0.42 | 0.03 | <0.001 | 0.42 | 0.03 | <0.001 |
| PVT, Dry Forest[2] | 0.26 | 0.03 | <0.001 | 0.26 | 0.03 | <0.001 |
| PVT, Other[2] | 1.39 | 0.11 | <0.001 | 1.39 | 0.11 | <0.001 |
| Step Length[1] | 0.04 | <0.001 | <0.001 | 0.04 | <0.001 | <0.001 |
| Turning Angle[3] | 0.04 | 0.01 | <0.001 | 0.04 | 0.01 | <0.001 |
| CP × D2C[1] | 0.46 | 0.22 | <0.001 | 0.45 | 0.13 | <0.001 |
| KP × D2K[1] | 0.08 | 0.10 | 0.43 | 0.01 | 0.10 | 0.95 |
| KP × D2K[1] × TSK[1] | -0.05 | 0.04 | 0.25 | -0.02 | 0.04 | 0.60 |

[1]Indicates the variable was natural log transformed; [2]the reference category was Grassland; [3]indicates the cosine of the variable was used.

**Table S8.** Parameter estimates for step selection functions for bobcats using different buffer sizes to determine cougar presence (CP). PVT = potential vegetation type, D2C = distance to nearest cougar, KP = kill present, D2K = distance to kill, and TSK = time since kill.

| | 500 m | | | 1500 m | | |
|---|---|---|---|---|---|---|
| | *β* | *SE* | *P* | *β* | *SE* | *P* |
| Canopy Cover | 0.32 | 0.01 | <0.001 | 0.32 | 0.01 | <0.001 |
| Distance to Road[1] | 0.10 | 0.02 | <0.001 | 0.10 | 0.02 | <0.001 |
| Distance to Water[1] | 0.01 | 0.02 | 0.45 | 0.01 | 0.02 | 0.46 |
| Ruggedness | 0.02 | 0.02 | 0.23 | 0.02 | 0.02 | 0.23 |
| PVT, Wet Forest[2] | 0.98 | 0.07 | <0.001 | 0.98 | 0.07 | <0.001 |
| PVT, Dry Forest[2] | 0.71 | 0.07 | <0.001 | 0.71 | 0.07 | <0.001 |
| PVT, Other[2] | 1.26 | 0.31 | <0.001 | 1.26 | 0.31 | <0.001 |
| Step Length[1] | 0.05 | 0.01 | <0.001 | 0.05 | 0.01 | <0.001 |
| Turning Angle[3] | 0.22 | 0.02 | <0.001 | 0.22 | 0.02 | <0.001 |
| CP × D2C[1] | 0.17 | 0.35 | 0.62 | -0.06 | 0.16 | 0.72 |
| KP × D2K[1] | 0.14 | 0.17 | 0.41 | 0.15 | 0.17 | 0.38 |
| KP × D2K[1] × TSK[1] | -0.09 | 0.07 | 0.19 | -0.10 | 0.07 | 0.17 |

[1]Indicates the variable was natural log transformed; [2]the reference category was Grassland; [3]indicates the cosine of the variable was used.

**Table S9.** Parameter estimates for step selection functions for coyotes using different buffer sizes to determine kill presence (KP). PVT = potential vegetation type, CP = cougar present, D2C = distance to nearest cougar, D2K = distance to kill, and TSK = time since kill.

| | 2,000 m | | | 4,000 m | | |
|---|---|---|---|---|---|---|
| | *β* | *SE* | *P* | *β* | *SE* | *P* |
| Canopy Cover | -0.13 | 0.01 | <0.001 | -0.13 | 0.01 | <0.001 |
| Distance to Road[1] | 0.09 | 0.01 | <0.001 | 0.09 | 0.01 | <0.001 |
| Distance to Water[1] | 0.02 | 0.01 | 0.01 | 0.02 | 0.01 | 0.01 |
| Ruggedness | -0.06 | 0.01 | <0.001 | -0.06 | 0.01 | <0.001 |
| PVT, Wet Forest[2] | -0.08 | 0.02 | <0.001 | -0.09 | 0.02 | <0.001 |
| PVT, Dry Forest[2] | -0.07 | 0.02 | <0.001 | -0.08 | 0.02 | <0.001 |
| PVT, Other[2] | -0.25 | 0.15 | 0.09 | -0.26 | 0.15 | 0.09 |
| Step Length[1] | 0.02 | <0.001 | <0.001 | 0.03 | <0.001 | <0.001 |
| Turning Angle[3] | -0.09 | 0.01 | <0.001 | -0.09 | 0.01 | <0.001 |
| CP × D2C[1] | 0.14 | 0.08 | 0.09 | 0.20 | 0.08 | 0.02 |
| KP × D2K[1] | -0.13 | 0.05 | 0.02 | -0.25 | 0.05 | <0.001 |
| KP × D2K[1] × TSK[1] | 0.05 | 0.02 | <0.001 | 0.06 | 0.01 | <0.001 |

[1]Indicates the variable was natural log transformed; [2]the reference category was Grassland; [3]indicates the cosine of the variable was used.

**Table S10.** Parameter estimates for step selection functions for black bears using different buffer sizes to determine kill presence (KP). PVT = potential vegetation type, CP = cougar present, D2C = distance to nearest cougar, D2K = distance to kill, and TSK = time since kill.

| | 2,000 m | | | 4,000 m | | |
|---|---|---|---|---|---|---|
| | *β* | *SE* | *P* | *β* | *SE* | *P* |
| Canopy Cover | 0.36 | 0.01 | <0.001 | 0.36 | 0.01 | <0.001 |
| Distance to Road[1] | 0.14 | 0.01 | <0.001 | 0.14 | 0.01 | <0.001 |
| Distance to Water[1] | -0.01 | 0.01 | 0.34 | -0.01 | 0.01 | 0.34 |
| Ruggedness | 0.16 | 0.01 | <0.001 | 0.16 | 0.01 | <0.001 |
| PVT, Wet Forest[2] | 0.42 | 0.03 | <0.001 | 0.42 | 0.03 | <0.001 |
| PVT, Dry Forest[2] | 0.26 | 0.03 | <0.001 | 0.26 | 0.03 | <0.001 |
| PVT, Other[2] | 1.39 | 0.11 | <0.001 | 1.39 | 0.11 | <0.001 |
| Step Length[1] | 0.04 | <0.001 | <0.001 | 0.04 | <0.001 | <0.001 |
| Turning Angle[3] | 0.04 | 0.01 | <0.001 | 0.04 | 0.01 | <0.001 |
| CP × D2C[1] | 0.51 | 0.15 | <0.001 | 0.54 | 0.15 | <0.001 |
| KP × D2K[1] | 0.12 | 0.09 | 0.17 | 0.03 | 0.07 | 0.68 |
| KP × D2K[1] × TSK[1] | -0.07 | 0.02 | <0.001 | -0.04 | 0.02 | 0.04 |

[1]Indicates the variable was natural log transformed; [2]the reference category was Grassland; [3]indicates the cosine of the variable was used.

**Table S11.** Parameter estimates for step selection functions for bobcats using two different buffer sizes to determine kill presence (KP). PVT = potential vegetation type, CP = cougar present, D2C = distance to nearest cougar, D2K = distance to kill, and TSK = time since kill.

| | 2,000 m | | | 4,000 m | | |
|---|---|---|---|---|---|---|
| | *β* | *SE* | *P* | *β* | *SE* | *P* |
| Canopy Cover | 0.32 | 0.01 | <0.001 | 0.32 | 0.01 | <0.001 |
| Distance to Road[1] | 0.10 | 0.02 | <0.001 | 0.10 | 0.02 | <0.001 |
| Distance to Water[1] | 0.01 | 0.02 | 0.46 | 0.01 | 0.02 | 0.45 |
| Ruggedness | 0.02 | 0.02 | 0.24 | 0.02 | 0.02 | 0.23 |
| PVT, Wet Forest[2] | 0.98 | 0.07 | <0.001 | 0.98 | 0.07 | <0.001 |
| PVT, Dry Forest[2] | 0.71 | 0.07 | <0.001 | 0.71 | 0.07 | <0.001 |
| PVT, Other[2] | 1.26 | 0.31 | <0.001 | 1.26 | 0.31 | <0.001 |
| Step Length[1] | 0.05 | 0.01 | <0.001 | 0.05 | 0.01 | <0.001 |
| Turning Angle[3] | 0.22 | 0.02 | <0.001 | 0.22 | 0.02 | <0.001 |
| CP × D2C[1] | 0.27 | 0.26 | 0.31 | 0.29 | 0.27 | 0.28 |
| KP × D2K[1] | 0.07 | 0.18 | 0.70 | 0.01 | 0.16 | 0.97 |
| KP × D2K[1] × TSK[1] | -0.08 | 0.06 | 0.16 | -0.04 | 0.05 | 0.48 |

[1]Indicates the variable was natural log transformed; [2]the reference category was Grassland; [3]indicates the cosine of the variable was used.

**Table S12.** Parameter estimates for a step-selection function for black bears separated by sex. PVT = potential vegetation type, CP = cougar present, D2C = distance to nearest cougar, KP = kill present, D2K = distance to kill, and TSK = time since kill.

| | Male | | | Female | | |
|---|---|---|---|---|---|---|
| | *β* | *SE* | *P* | *β* | *SE* | *P* |
| Canopy Cover | 0.35 | 0.01 | <0.001 | 0.38 | 0.01 | <0.001 |
| Distance to Road[1] | 0.09 | 0.02 | <0.001 | 0.25 | 0.03 | <0.001 |
| Distance to Water[1] | -0.04 | 0.01 | <0.001 | 0.03 | 0.01 | 0.03 |
| Ruggedness | 0.15 | 0.01 | <0.001 | 0.17 | 0.01 | <0.001 |
| PVT, Wet Forest[2] | 0.43 | 0.04 | <0.001 | 0.45 | 0.06 | <0.001 |
| PVT, Dry Forest[2] | 0.22 | 0.04 | <0.001 | 0.35 | 0.05 | <0.001 |
| PVT, Other[2] | 0.60 | 0.17 | <0.001 | 2.37 | 0.16 | <0.001 |
| Step Length[1] | 0.03 | 0.01 | 0.09 | 0.06 | 0.02 | <0.001 |
| Turning Angle[3] | 0.03 | <0.001 | <0.001 | 0.05 | 0.01 | <0.001 |
| CP × D2C[1] | 0.63 | 0.19 | <0.001 | 0.48 | 0.25 | 0.05 |
| KP × D2K[1] | -0.13 | 0.12 | 0.30 | 0.25 | 0.16 | 0.13 |
| KP × D2K[1] × TSK[1] | 0.04 | 0.05 | 0.40 | -0.12 | 0.07 | 0.06 |

[1]Indicates the variable was natural log transformed; [2]the reference category was Grassland; [3]indicates the cosine of the variable was used.

**Movie S1 (separate file).** Video clip from remote camera deployed at an elk carcass killed by a GPS-collared cougar. The cougar chases off three coyotes which can be heard vocalizing in the background. This clip occurred immediately after the photograph in Fig. 1b was taken.

**Movie S2 (separate file).** Video clip from remote camera deployed at an elk carcass killed by a GPS-collared cougar. A coyote can be seen and heard bark-howling in the upper left as the cougar feeds in the bottom right of the frame.

**Movie S3 (separate file).** Video clip from remote camera deployed at an elk carcass killed by a GPS-collared cougar. A coyote can be seen and heard bark-howling in the upper left as the cougar vocalizes and then leaves the carcass. The coyote then approaches the carcass to feed.
